# Resident memory T cell precursors in tumor draining lymph nodes require type-1 IFN for optimal differentiation

**DOI:** 10.1101/2023.08.02.551658

**Authors:** Nikhil Khatwani, Tyler Searles, Jichang Han, Cameron Messier, Neeti Mittal, Aaron Hawkes, Aleksey K. Molodstov, Delaney E. Ramirez, Owen Wilkins, Yina H. Huang, Fred W. Kolling, Pamela C Rosato, Mary Jo Turk

## Abstract

Resident memory (Trm) cells play an essential role in anti-tumor immunity. However, little is known about the precursors that differentiate into protective Trm populations against cancer. Here we employed an established model of B16 melanoma neoadjuvant anti-CD4 therapy, to track tumor antigen-specific CD8+ T cells through tissues and across time; from their priming as effectors to their differentiation into Trm. We show that tumor-draining lymph nodes (TDLNs) contain Teff cells that begin to express canonical Trm markers CD103 and CD69. These tumor-specific Teff cells seeded skin and tumor during the effector phase of the response, although egress from these tissues was not required Trm development in LNs. Paired scRNAseq/scTCRseq was used to identify Teff clonotypes in TDLNs and trace their differentiation, in real-time, into Trm populations. We found that expanded clonotypes favored the Trm fate and were unlikely to co-differentiate into other lineages. Precursors of Trm (pre-Trm) clonotypes that subsequently seeded populations throughout tumors, LNs, and skin, were characterized by early expression of tissue residency, stemness, and type-1 IFN sensing genes. These multipotent pre-Trm cells sensed plasmacytoid dendritic cell-derived type-1 interferons in TDLNs, and their expression of interferon alpha receptor was required for their formation of Trm populations in LNs but not in skin. These findings reveal the defining features of pre-Trm cells in response to tumor antigens, and reveal a previously unappreciated role for type-1 IFNs in programming regional Trm immunity to cancer.

**One Sentence Summary:** Anti-tumor effector CD8 T cells adopt early characteristics of tissue residency and stemness, and rely on the sensing of type-1 interferons for their local differentiation into resident memory T cells.

## INTRODUCTION

Cancer is a complex and devastating disease characterized by uncontrolled cell growth and subsequent metastasis (*1, 2*). Despite significant advances in treatment, metastasis continues to be a leading cause of cancer morbidity and mortality (*1*). The immune system plays an important role in tumor prevention (*3, 4*), and T cells are critical in mediating effective anti-tumor immune responses (*5*). One of the primary objectives of cancer immunotherapy is to establish immunological memory for sustained tumor suppression by means of T cells that persist for years in visceral organs where tumors may recur or metastasize (*4, 6–9*).

The immune system’s memory for a previously encountered antigen is one of its most essential properties (*4, 10*). Upon encounter with their cognate antigen, naive T cells undergo clonal expansion and develop into effector T cells. These clonally expanded cells then endure a bifurcation in differentiation, producing short-lived terminally differentiated effector T cells and memory precursor T cells (*11*). Subsequent generation of memory T cells can be broadly classified by their circulating or tissue resident characteristics. Circulating memory, T_CIRM_ is comprised of central memory (T_CM_) and effector memory (T_EM_) cell subsets. T_CM_ cells express lymphoid homing molecules CD62L and CCR7, whereas T_EM_ cells expresses neither (*12*). Tissue-resident memory (T_RM_) cells, unlike their T_CIRM_ counterparts, lack circulating molecules but express tissue retention molecules such as CD103 and CD69 (*12–14*). Trm cells demonstrate prompt activation (*15*) and proliferation upon antigen re-encounter (*16*), and are thereby positioned as first responders during tumor recurrence. As such, a growing body of literature supports the perception that T_RM_ cells can be more potent than T_CIRCM_ cells in mediating the anti-tumor response (*9, 17–23*).

T_RM_ cells have been identified in a variety of tumors, including melanoma, lung, breast, colon, and ovarian cancer, and more recently, at metastatic sites, in association with improved overall survival (*7, 24, 25*). However, generating a stable pool of widely disseminated tumor-specific T_RM_ cells necessitates a thorough understanding of pathways driving their programming and differentiation (*26*). Studies have characterized T_RM_ precursor cells exhibiting a distinct transcriptome imprinted after entry into peripheral tissues, allowing for clear distinction from their circulating counterparts (*27–30*). However, recent reports have identified effector T cells expressing canonical T_RM_ markers CD69 and CD103 that are superior in giving rise to T_RM_ cells (*31, 32*) with a propensity that was acquired clonally in the circulating effector pool, even prior to tissue entry (*33*). Interestingly, in certain contexts, T_RM_ predisposition may occur even at the naïve T cell stage (*34, 35*) and can be regulated by signals derived from antigen presenting cells (*36–38*). Furthermore, transcription factors, including *Hobit, Blimp1, Runx3* and *mTOR* have been identified in mouse studies to promote early T_RM_ skewing (*27, 29, 30, 33, 39, 40*), by balancing residency and circulating molecules *Cd103, Cd69, S1pr1, Sell* (*27–29, 41–43*). However, the majority of this understanding of T_RM_ precursors derives from studies in viral models, while little is known about when, where, and how T_RM_ cells develop in the context of cancer.

The present studies were based on a hypothesis that broadly disseminated tissue residency can be established by T_RM_ precursors early during an anti-tumor response, and that such tissue specific T_RM_ fate decisions are governed by exposure to precise molecular signals during priming. Here, in an established model of melanoma Ag-specific Trm generation following therapeutic neoadjuvant anti-CD4 regulatory T cell (Treg) depletion, we employ locational tracking and clonal mapping of tumor antigen specific CD8+ T cells during priming and memory establishment, to identify features that demarcate transcriptionally distinct T_RM_ precursor cells. Our findings reveal a crucial role for early type 1 interferon sensing by T_RM_ precursors in promoting their tissue distribution and subsequent T_RM_ establishment, specifically in lymph nodes. These studies support a role for factors within tumor draining lymph nodes (TDLNs) that can instruct the retention and programming of regionally protective T_RM_ responses.

## RESULTS

### A subset of tumor-specific CD8 effector T cells acquires features of residency and stemness in tumor-draining lymph nodes

To identify T_RM_ precursor populations in the setting of anti-tumor immunity, we utilized our established model of B16 melanoma neoadjuvant treatment, wherein administration of anti-CD4 monoclonal antibody to deplete Tregs, followed by surgical tumor excision, results in autoimmune vitiligo and robust T_RM_ formation throughout skin and draining lymph nodes (*23, 44*). These Trm populations are durably protective against melanoma rechallenge in skin and in inguinal TDLNs (*23, 44, 45*). We previously showed that these Trm populations recognize shared melanoma/melanocyte antigens, and that they can be tracked using sentinel gp100_25-33_-specific pmel T cells (*23, 44, 45*). We also showed that pmel cells harvested from tumor-bearing mice on day 12 could confer Trm populations to the skin of recipient mice following adoptive transfer (*44*), suggesting the presence of early Trm precursors in TDLNs. However, we had yet to understand the properties that defined these precursors or the factors that determined their fate as resident memory.

To understand which tumor-specific effector (Teff) cells serve as precursors to Trm, we performed single cell (sc)RNA sequencing on CD44^+^ pmel cells sorted from day 12 TDLNs. By co-clustering with our previously published dataset of pmel cells at a memory timepoint (day 45) from LNs and skin, we sought to identify a day 12 population that transcriptionally resembled Trm cells (Fig 1A). Gene expression analysis on 1961 sorted pmel T cells revealed that clustering was determined largely by timepoint. Whereas day 45 pmel populations previously clustered based on tissue of occupancy (*23*) inclusion of day 12 cells drove skin and LN memory T cells together into two main clusters (C0 and C5), with day 12 cells distinctly forming their own clusters (Pmel_C1 and Pmel_C2) (Fig 1B&C). Type-1 IFN sensing Trm and stem-like memory LN sub-populations were still identified in LNs as we previously published (*23*), but these clusters were also distal to the day 12 clusters (Fig. 1C-E).

**Fig. 1.**
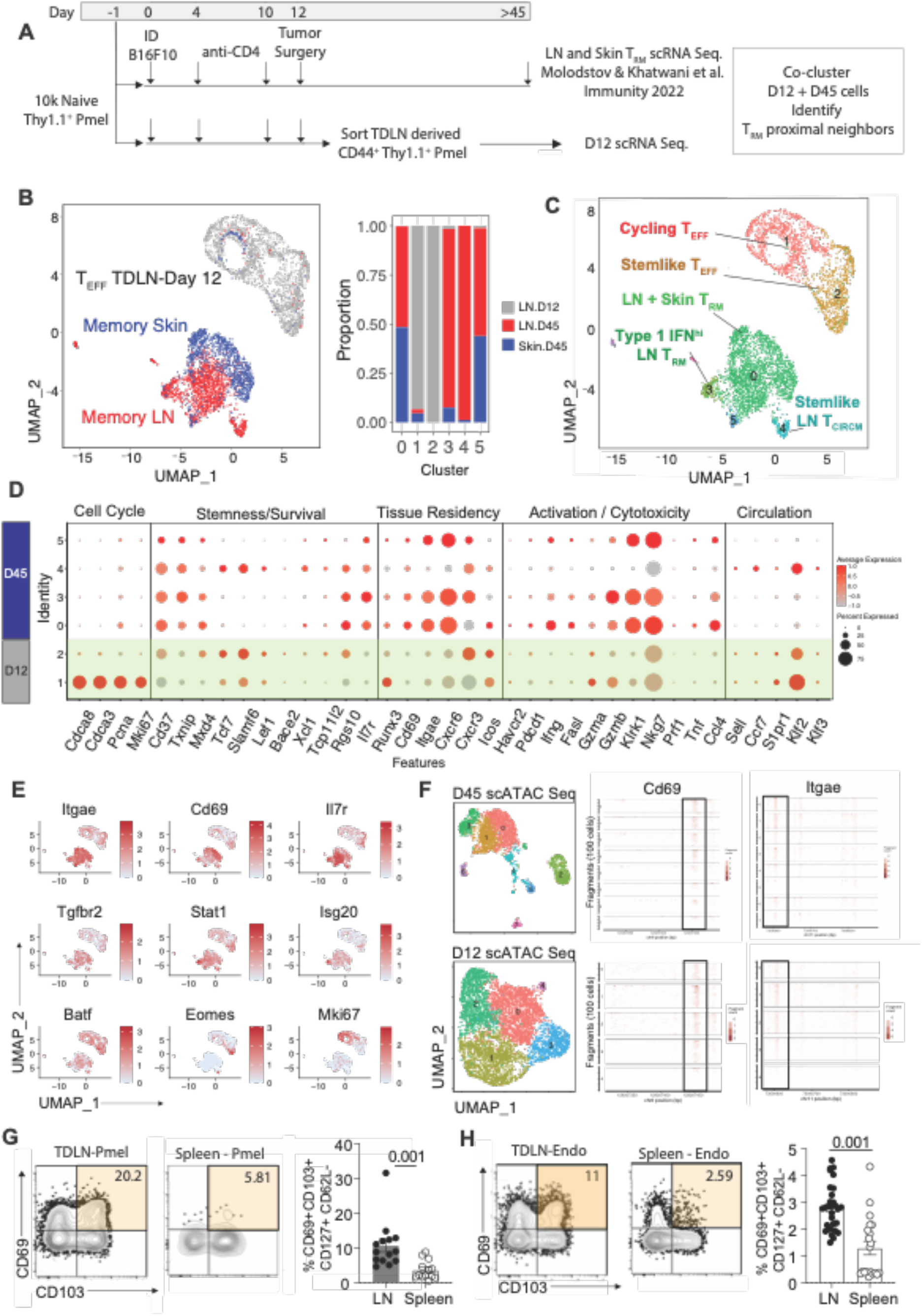
Trm transcriptional, epigenetic, and phenotypic features appear on tumor Ag-specific Teff cells in TDLNs. (**A**) Schematic showing B16 melanoma neoadjuvant treatment model. Pmel cells were FACS sorted from tumor-draining lymph nodes (TDLN) of n=5-8 mice, pooled, on the day of surgery (d12; priming). (**B-left**) UMAP plot displaying single cell (sc) RNA sequencing results from 1961 sorted TDLN pmel cells, memory LN and vitiligo skin (published dataset). (**B-right**) Proportion of cells present in individual clusters corresponding to tissues (clusters with fewer than 50 cells (C6-7) were omitted from further analysis). (**C**) UMAP plot with clusters labeled based on their transcriptional features. (**D**) Dot plot of gene expression from scRNA sequencing on pmel cells (**E**) Feature plot of genes associated with residency, stemness and type 1 IFN sensing. (**F**) UMAP plots with scATAC sequencing results on pmel cells sorted from TDLN, d45-top and d12-bottom. Chromatin accessibility Tileplots are shown for *Cd69* and Cd103. (**G-H**) Flow cytometry phenotype analysis of pmel cells (**G**) and polyclonal CD8+ T cells (**H**) harvested from TDLN and spleen of mice treated as in panel A, on d12. Bar graphs with proportions of indicated phenotypes. Symbols represent individual mice; horizontal lines depict means; significance was determined by unpaired t test, with n.s. (non-significant) denoting p > 0.05. Flow plots depict representative data from two to three independent experiments, with n=10-30 mice, 5-8 per group; mean ± SEM.

Of the two day 12 Teff clusters, Pmel_C1 displayed stronger features associated with effector function (*Gzma, Gzmb, Fasl)* and cell cycle/proliferation (*Cdca8, Cdca3, Ki67*), whereas Pmel C2 was lower for cell cycle genes, and instead expressed markers associated with memory (*Eomes, Cd127), and* stemness (*Tcf7, Slamf6)*. Pmel_C2 also expressed some genes associated with tissue residency (*Cxcr3, Icos, Runx3, Tgfbr2, Batf*) (*11, 27, 29, 46–48*) (Fig 1D), however, both day 12 clusters lacked strong expression of classical Trm transcripts *Cd103 and Cd69* (Fig 1E), particularly in comparison to the day 45 Trm clusters.

To investigate if day 12 pmel Teff cells had evidence of chromatin accessibility in Trm-associated *Cd69 and Cd103* gene loci, we separately performed scATAC sequencing. Interestingly, at least three day 12 pmel clusters had open chromatin in *Cd69* and *Itgae* loci, at levels comparable to the day 45 memory populations (Fig 1F). To determine if these Trm markers were actually expressed on Teff cells, flow cytometry was performed. Indeed, ∼20% of pmel Teff cells in TDLNs exhibited a Trm-like (CD69^+^CD103^+^) phenotype, and this population was absent from the spleen where Trm population do not form (*23*) (Fig 1G). Cells with a CD62L^low^CD127^hi^CD69^+^CD103^+^ Trm-like phenotype accounted ∼10% of the pmel population and ∼3% of the endogenous CD8^+^CD44^+^ Teff population in TDLNs on day 12 (Fig. 1G-H, Fig. S1A). As these data suggested that a Trm-like sub-population was being missed in our day 12 scRNAseq analysis, we separately clustered only day 12 cells (without day 45), which indeed resolved two smaller sub-clusters with more pronounced *Cd69* and *Itgae* expression, additional Trm genes and several stemness markers (Fig. S1B-D). Overall, these data revealed the presence of a tumor-specific Teff cell subpopulation in TDLNs that adopts features of tissue residency and stemness prior to Trm differentiation in TDLNs.

### Trm differentiation in TDLNs does not require retrograde migration from tissue

Studies in viral infection models have shown that retrograde migration of T cells from skin and lung generates T_RM_ populations in tissue-draining LNs (*49, 50*). To understand how early Trm features are established in TDLNs, we next investigated if pmel cells underwent retrograde migration from skin and tumors to TDLNs during the effector phase of the response. We tracked migration of tumor-specific T cells using pmel cells expressing the photoconvertible Kaede marker which is converted from GFP to RFP *in situ* by UV exposure (*51*). Indeed, UV photoconversion of dermal B16 tumors resulted in efficient (>80%) T cell photoconversion from GFP to RFP in tumors without leakiness in the TDLN (Fig. S2A). To capture pmel T cell egress to TDLNs during the peak of the response (*52*), tumors were photoconverted on days 10 and 11, and LNs were evaluated one day later (Fig. 2A). Indeed, tumor-derived (RFP+) pmel cells were detected in the TDLN on day 12, although this population was small, accounting for only 1-3% of the pmel population in LNs (Fig. 2B-C). Additionally, whereas LNs contained a population of CD103^+^CD69^+^ pmel cells, CD69+ pmel cells in tumors lacked CD103, and those migrating from tumors to TDLNs failed to acquire CD103 (Fig. 2B, D). Rather, tumor-derived pmel cells tended to express more PD-1 as compared with their tumor non-derived counterparts in LNs (Fig. 2D, E). Therefore, retrograde migration from tumor was not pronounced, nor was it associated with acquisition of Trm features in the TDLN.

**Figure 2:**
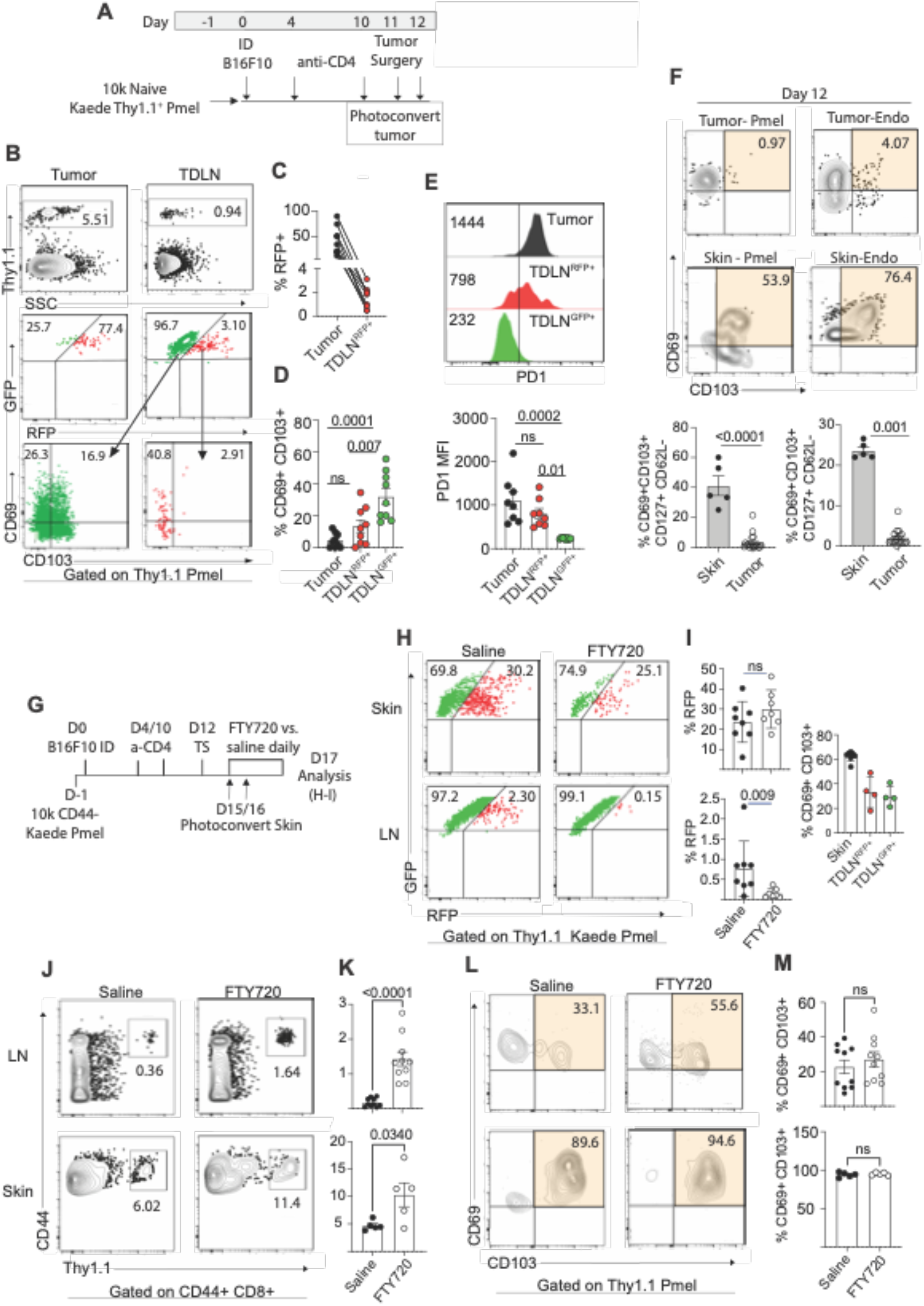
R**e**trograde **migration of precursors from tumor or skin to draining lymph nodes is not required for Trm differentiation.** (**A**) Schematic showing tumor photoconversion using Kaede pmel T cells. (**B**) (**Top**) Proportion of CD8^+^CD44^+^ pmel cells on d12. (**B-Middle, C**) Proportion of Thy1.1^+^GFP^+^ or Thy1.1^+^ GFP^+^ RFP^+^ (photoconverted) pmel cells. (**B-Bottom, D**) Proportion of CD69^+/-^CD103^+/-^ pmel cells gated on CD8^+^CD44^+^ Thy1.1^+^GFP^+^ vs. GFP^+^RFP^+^. (**E**) PD1 geometric mean for CD8^+^CD44^+^Thy1.1^+^GFP^+^ vs. GFP^+^RFP^+^ pmel cells. (**F**) Proportion of CD69^+/-^ CD103^+/-^ gated on CD8^+^ CD44^+^ Thy1.1^+^ pmel cells (**left**) or CD8^+^ CD44^+^ open repertoire (**right**) in tumor bearing mice on d12. (**G**) Schematic showing skin photoconversion. (**H**) proportion of GFP^+^ vs. GFP^+^RFP^+^ (photoconverted) cells in each tissue five days after tumor excision (d12+5). (**I**) proportion of GFP^+^ RFP^+^ cells (**left**) and CD69^+^ CD103^+^ pmel cells (**right**). (**J-K**) proportion of CD44+ pmel cells and (**L-M**) CD69^+/-^ CD103^+/-^ 30-60 days after tumor surgery (memory timepoint). Symbols represent individual mice; horizontal lines depict means; significance was determined by paired (**C-E**) or unpaired t test (**F**), with n.s. (non-significant) denoting p > 0.05. Flow plots depict representative data from two independent experiments; n=5-8 mice/group; mean ± SEM (**A-F, J-M**) or taken from a single experiment; n=6-9; mean ± S.D (**H-I**).

Based on the literature, it remained likely that skin was a source of Trm cells in LNs. Indeed, in contrast to the tumor, pmel T cells (and endogenous CD8 T cells) in the skin already exhibited a pronounced CD62L^low^CD69^hi^CD103^hi^CD127^hi^ Trm phenotype on day 12 (Fig. 2F, Fig. S2B). Because T cell population in the skin are difficult to quantify at early time points (*44*), we assessed migration from skin surrounding the surgical site during a 5-day window immediately following tumor excision (Fig. 2G). Indeed, we observed skin-derived RFP+ pmel cells migrating to LNs during this time. However, as with tumor, the egressed pmel cell population accounted for only a small proportion (∼1-2%) of pmel cells in TDLNs (Fig 2H). Of note, skin-egressed cells were more likely to exhibit a CD69^+^CD103^+^ phenotype (Fig. 2I, Fig S2C-D) than what had been observed with tumor. Because of this, we reasoned that continual Trm cell egress from skin, particularly during the development of autoimmune vitiligo, could gradually establish the Trm response in TDLNs. To test this hypothesis, we employed the S1PR1 antagonist FTY720, which inhibits lymphocyte egress from secondary lymphoid organs and peripheral tissues (*53, 54*) and which we found efficiently blocked Kaede-pmel T cell egress from skin to TDLNs (Fig 2H-I). Mice were treated with FTY720 starting one day after tumor excision and continuing for 45 days to establish memory. However, contrary to our hypothesis, proportions of pmel cells in both TDLN and skin were substantially augmented by FTY720 treatment (Fig 1 J, K). Pmel cells in LNs and skin of FTY720-treated mice were also unimpaired in their acquisition of a CD69^hi^CD103^hi^ phenotype (Fig. 1L, M) and expression of Trm markers CXCR6 and CD127 (Fig. S2E, Fig. 1D). To confirm that FTY720 was not locking T cell populations in tissue that weren’t truly resident there, we separately treated with FTY720 for 1 month and then discontinued treatment for 2 weeks before analysis (Fig. S2F). In this setting FTY720 still augmented proportions of pmel cells with a Trm phenotype in both skin and LNs (Fig. S2 G-H). Thus, contrary to prior work in viral models (*49, 50*), tumor-specific T_RM_ differentiation proceeded locally and discretely in skin and LNs without a requirement for retrograde migration. These data also establish that tumor-specific Trm precursors (pre-Trm) seed their tissues of residency by day 12 of the response.

### Trm precursors adopt a resident/stem-like transcriptional profile in TDLNs

Analysis of day 12 pmel cells revealed Teff cells in TDLNs that acquire early features of stemness and residency. However, conclusions generated with pmel cells may not fully recapitulate the endogenous CD8 T cell repertoire that constitutes protective Trm in this model (*23, 44*). Thus, we next sought to define Trm precursors within the endogenous CD8 Teff cell compartment in TDLNs. To this end we conducted scRNA sequencing on antigen-experienced CD44^+^ CD8^+^ T cells sorted from day 12 surgically excised TDLNs (Fig 3A).

**Figure 3:**
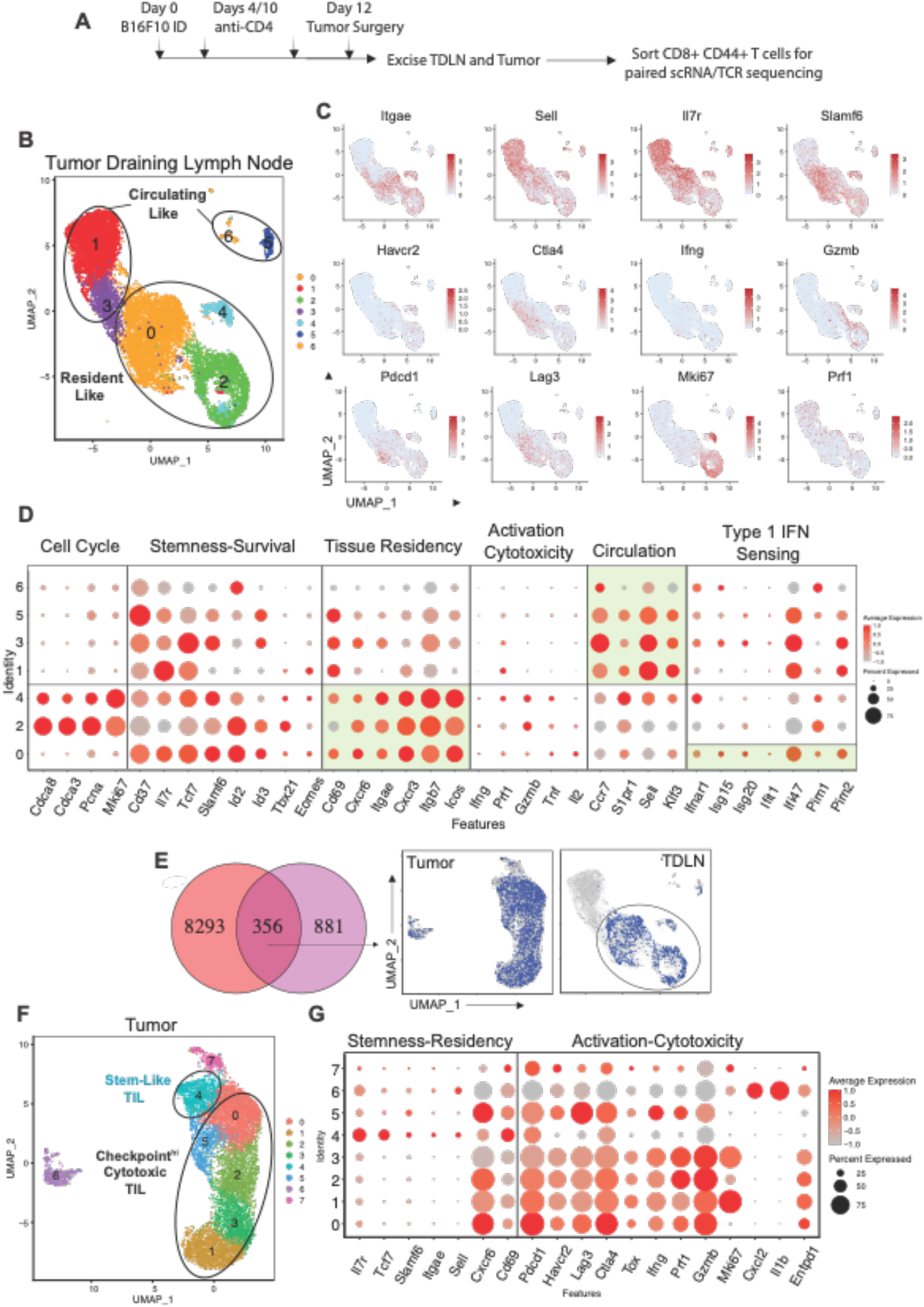
E**n**dogenous **CD8 T cells acquire Trm and stemness traits in TDLN in conjunction with type 1 interferon sensing.** (**A**) Schematic showing B16 melanoma neoadjuvant treatment model. Endogenous polyclonal CD8^+^ CD44^+^ T cells were FACS sorted from tumor-draining lymph node (TDLN) and tumor (n=3) on the day of surgery – d12 (priming). Sorted cells were submitted for paired scRNA/TCR sequencing. (**B**) scRNA UMAP plots displaying 12,377 TDLN derived CD8^+^ CD44^+^ T cells with clusters labeled based on their transcriptional profiles. (**C**) Feature plots on expression of selected genes on d12 TDLN CD8^+^ T cells. (**D**) Dot plot depicting gene expression of markers associated with cell cycle, stemness/survival, residency, activation, circulation, and type-1 IFN sensing. (**E**) (**Left**) Venn-diagram showing clonal overlap of CD8^+^ CD44^+^ T cells between TDLN and tumor. (**Right**) scRNA UMAP plots displaying matched clones on each tissue of origin. (**F**) scRNA UMAP plots displaying 10,000 tumor-derived CD8^+^ CD44^+^ T cells with clusters labeled based on their transcriptional profile. (**G**) Dot plot depicting gene expression of markers associated with cell cycle, stemness/survival, residency, activation, and circulation from tumor derived CD8^+^ CD44^+^ T cells. (**B, C, D**) Results are representative of two independent scRNA sequencing experiments, with n=3 pooled mice in each experiment.

Transcriptional profiling revealed the presence of seven distinct Teff clusters which could be broadly subclassified by their dichotomous expression of residency vs. recirculation-associated genes (Fig. 3B). Clusters TDLN_C0, C2, and C4 were termed “Resident-like” based on their expression of core Trm-associated genes (e.g. *Itgae, Cxcr6, Cxcr3, Itgb7* and *Icos*) and weak expression of circulating memory (Tcirm) markers (e.g. *Ccr7, S1pr1, Klf3* and *Sell)*, whereas clusters TDLN_C1, C3, C5, and C6 were termed “Circulating-like” based on their reciprocal lower expression of Trm genes and higher expression of Tcirm markers (Fig. 3B-D, Fig. S3A). An exception was *Cd69*, which was broadly expressed across the clusters presumably marking activation (Fig. 3D). Closer examination of the resident-like clusters showed that TDLN_C2 and TDLN_C4 also had strong expression of cell-cycle/proliferation associated transcripts (*Cdca8, Cdca3, Pcna and Mki67)*, whereas TDLN_C0 lacked these (Fig. 3C, D). All three resident-like clusters expressed transcripts associated with stemness/pro-survival (e.g., *Cd37, Il7r, Tcf7, Slamf6, Id2* and *Id3*), but this was most prominent in TDLN_C0, which interestingly also expressed genes associated with type-1 IFN signaling (Fig. 3C, D). Thus, as observed for pmel cells (Fig. 1), endogenous Ag-experienced CD8 Teff cells in the TDLN also adopt transcriptional signatures indicative of tissue residency and stemness.

Another major benefit to tracking the endogenous repertoire was the ability to track individual T cell clonotypes. To gain an understanding of tumor Ag-specificity within the TDLN compartment, we also performed scTCR sequencing on these day 12 Teff cells and defined clonal overlap with CD8^+^CD44^+^ T cells from surgically excised B16 tumors harvested on the same day (Fig 3A). Indeed, 356 clonotypes shared occupancy in both tissues (Fig 3E). Interestingly, essentially all tumor-matched clonotypes were confined to the three resident-like clusters (TDLN_ C0, C2, and C4) in TDLNs (Fig 3E), which were more clonally expanded than the circulating-like clusters (Fig. S3B). Taken with our pmel data, this supports the conclusion that tumor Ag-specific CD8 Teff cells which were clonally expanding were preferentially adopting features of residency and stemness in TDLNs.

We also performed scRNAseq on the CD8 T cells from excised tumors (TILs) to define the transcriptional features of TDLN-matched clonotypes in tumors. TILs formed eight clusters, the majority of which expressed markers associated with activation/exhaustion (e.g. *Pdcd1, Tim3, Ctla4* and *Lag3*) and function/cytotoxicity (e.g. *Ifng, Gzmb, Prf1,* and *Tnf*) (Fig. 3F, G). Some tissue residency features were apparent on CD8 T cells in the tumor (e.g. high *Cd69* and *Cxcr6*; low *Sell*) although others were absent (e.g. *Itgae* and *Il7r* expression). This is consistent with our flow cytometry data (Fig. 2) and supports the conclusion that Trm populations were absent from tumors (Fig. 3G, Fig. S3C, Fig 2F), although day 12 was understandably too early for memory differentiation. Interestingly, one TIL cluster (TIL_C4) expressed more of a stem-like signature (high *Tcf7 and Il7r,* low activation/cytotoxicity) (Fig. 3F, G) and was less clonally expanded than the other clusters (Fig. S3D). A look back at clonal matches however, showed strongest overlap of TDLN clonotypes across the activated/cytotoxic TIL clusters (TIL_C0, C1, C2, and C5), with less apparent overlap in the stem-like TIL_C4 (Fig. 3E). Thus, taken with data from our migration studies (Fig. 2), these findings support the interpretation that clonally expanding Teff cells in TDLNs simultaneously seed cytotoxic Teff cells in tumors and Trm precursors in skin, while also serving as precursors for Trm differentiation in LNs.

### Skin and LN Trm cells originate from stemlike-Trm precursors in TDLNs

A final benefit to tracking the open repertoire of CD8 Teff cells was the ability to follow the fate of putative Trm precursor clonotypes in real-time, i.e. in the same mice whose tumors and TDLNs had been excised and analyzed on day 12. Our prior work has shown that, in addition to the TDLNs, robust pmel Trm populations also develop in axillary and brachial LNs (*23*). Moreover, we conducted separate scTCRseq analyses which confirmed robust clonal intermixing between TDLNs, contralateral LNs and tumors prior to day 12 (Fig. S4A, B). Thus, we reasoned that Teff clonotypes from surgically excised TDLNs on day 12 would be maintained in vitiligo-affected skin and in axillary LNs as memory. Considering that individual mouse clonal repertoires are private (Fig. S4C), we survived the mice whose tumors and TDLNs had been analyzed in Fig. 3 and analyzed their skin and LNs forty five days after surgery.

To ensure enrichment of Trm cells from these specimens, we sorted CD8^+^CD44^+^T cell populations that were low/intermediate for CD62L (Fig. 4A, Fig. S5A). From skin, 1288 T cells formed 9 clusters (Fig. 4B), all of which expressed canonical tissue residency markers (*Itgae, Cd69, Il7r, Cxcr6,* and *Btg1*), and lacked tissue egress genes (*Sell, Ccr7*) (Fig 4C) (*23, 29, 55–59*). This was in accordance with sorted pmel Trm cells from skin (Fig. 1) and confirmed that essentially all CD8 T cells from vitiligo-affected skin adopt a Trm fate. However, there was some heterogeneity in the repertoire, e.g. with Skin_C2 having lower expression of certain Trm genes (*Nr4a1, Vps37b, and Icos)*, Skin_C3 expressing higher cytotoxicity-associated transcripts (*Gzmc and Ifitm2*) (*60–62*), and Skin_C6 having evidence of more robust TGF-β signaling (highest *Btg1* and *Malat1*) (*63*) (Fig 4C, Fig. S5B). In contrast, cells sorted from LNs were far more heterogeneous, with 12,377 T cells forming 10 clusters, most of which expressed transcripts associated with Tcirm (high *Ccr7, Cd62l, S1pr1, Cd127*) (Fig 4D, Fig. S5C) (*64*). However, one cluster (LN_C5) emerged as having low Tcirm transcripts and high expression of canonical Trm genes (e.g. *Itgae, Cxcr6, Il7r,* and *Cd69),* consistent with Trm (Fig. 4E). GSEA analysis further confirmed positive enrichment of a core LN T_RM_ signature in LN_C5 (*23*) (Fig 4F). Consistent with the protective function of Trm cells in LNs (*23*) LN_C5 also expressed effector-associated transcripts (*Gzmb, Ifng, Prf1*, and *Tnfa*) and had low levels of exhaustion-related transcripts (e.g. *Tox, Pdcd1, Tigit, Lag3* and *Havcr2*) (Fig. 4E, Fig. S5C). On the other hand, LN_C3 expressed high levels of *Tox, Pdcd1, Tigit*, Lag3, *Havcr2* and *Eomes,* consistent with classification as a progenitor/exhausted cluster (Fig. 4E, Fig. S5C-D). (*65*) (*66*). Thus, an array of endogenous skin and LN Trm populations were readily apparent on day 45, serving as a backdrop on which clonal matches to day 12 could be investigated.

**Figure 4:**
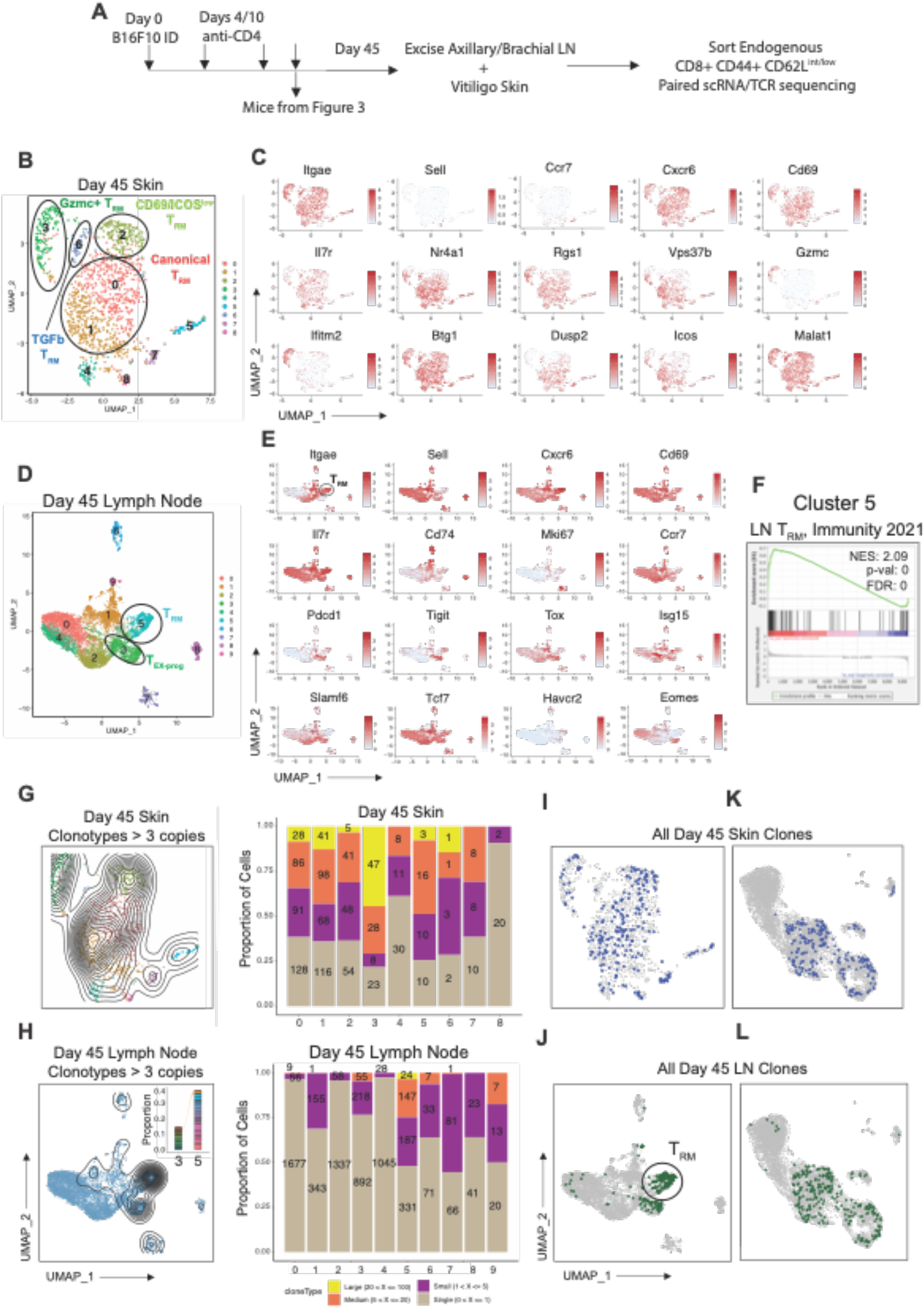
F**u**lly **differentiated Trm cells in skin and LNs have high clonal overlap with resident-like Teff cells from day 12.** (**A**) Schematic showing how endogenous polyclonal CD8^+^ CD44^+^ CD62L^low-int^ T cells were FACS sorted from axillary-brachial LNs, and CD8+ CD44+ cells were sored from vitiligo-affected skin on day 45 (d45); using the same mice that previously underwent analyses on d12 (in Fig. 3A; n=3). Sorted cells were submitted for paired scRNA/scTCR sequencing. (**B**) scRNA UMAP plot displaying 1288 CD8+ T cells from d45 skin, with clusters labeled based on their transcriptional profiles. (**C**) Feature plots of gene expression in skin CD8 T cells. (**D**) scRNA UMAP plot displaying 9400 d45 LN CD8+ T cells, with clusters labeled based on their transcriptional profiles. (**E**) Feature plots of gene expression in d45 LN CD8 T cells. (**F**) Gene set enrichment score analysis on T_RM_ cluster 5 from d45 LNs. Reference dataset used is from Molodtsov and Khatwani et al. 2021 LN T_RM_ gene signature. (**G, H**) **Left**-scRNA UMAP plots displaying clonal density for clonotypes > 3 copies. **Right**-scTCR data displaying proportion of clonal expansion spanning individual clusters from scRNA analysis. (**I-J**) D45 skin (**I**) and d45 LN (**J**) scRNA UMAP plots highlighting all TDLN matched clonotypes. (**K-L**) D12 TDLN scRNA UMAP highlighting all d45 skin (**K**) or lymph node (**L**) matched clonotypes.

Indeed, paired scTCRseq analysis revealed high clonal expansion throughout the skin Trm clusters (Fig. 4G), with expanded clonotypes also occupying multiple clusters with no substantial exclusivity to any particular cluster (Fig. S5E). This is consistent with the focused recognition of shared melanoma/melanocyte Ags by Trm cells in vitiligo-affected skin (*44*). Interestingly, whereas far less clonal expansion was detected in LNs overall, the Trm cluster (LN_C5) stood out as the most clonal (Fig 4H), illustrating that Trm cells became the most expanded memory population in LNs. The progenitor/exhausted cluster (LN_C3) contained moderately expanded clonotypes, although these same clonotypes did not co-occur in the Trm cluster (Fig. 4H inset, Fig. S5F). Thus, Trm and progenitor/exhausted populations in LNs appeared to be generated as discrete lineages.

Finally, we sought to determine whether clonal counterparts of the skin and LN memory populations on day 45 were represented earlier within Teff cells from the TDLNs. We also wished to understand the transcriptional features of such pre-Trm cells. Indeed, we found day 12 clonal matches in both the skin and LNs at day 45 (Fig. 4I). These persisting clonotypes were distributed across the Trm clusters in skin, and in LNs they were most pronounced in the Trm cluster as well as the progenitor/exhausted cluster (Fig. 4J). To ensure that we were not missing persistent Tcirm populations, we separately performed scTCRseq clonal tracing without CD62L^low^ selection/Trm enrichment, and here we found no persisting LN clonotypes from Day 12 (Fig. S6). Thus, Trm cells were the dominant memory populations generated. Importantly, retrospective identification of persisting/ memory clonotypes on the day 12 UMAP plots (from Figure 3) revealed that pre-Trm cells were segregated to the three resident/stem-like Teff clusters, with essentially no representation in recirculating Teff clusters (Fig. 4K-L). Taken together, these data confirmed the origination of Trm cells from resident/stem-like Teff cells in TDLNs, which were destined to become Trm without taking on other memory lineages.

### Type-1 IFN signals are required for a multipotent Trm fate

Having established the presence of Trm precursors in TDLN, we next sought to define the behavior of individual clonotypes that were destined for the Trm fate. To do this we generated enrichment scores for each of the pre-Trm clonotypes across each of the day 12 TDLN clusters. Interestingly, of 85 pre-Trm clonotypes that became Trm in skin, the vast majority had representation in the resident/stemlike TDLN_C0, and only some had co-representation in the other two resident-like clusters (TDLN_C2 and C4) (Fig. 5A). A similar pattern was observed for the 36 clonotypes that became Trm in the LNs, with dominant representation in TDLN_C0, and far less occurrence in the other clusters (Fig. 5B).

**Figure 5:**
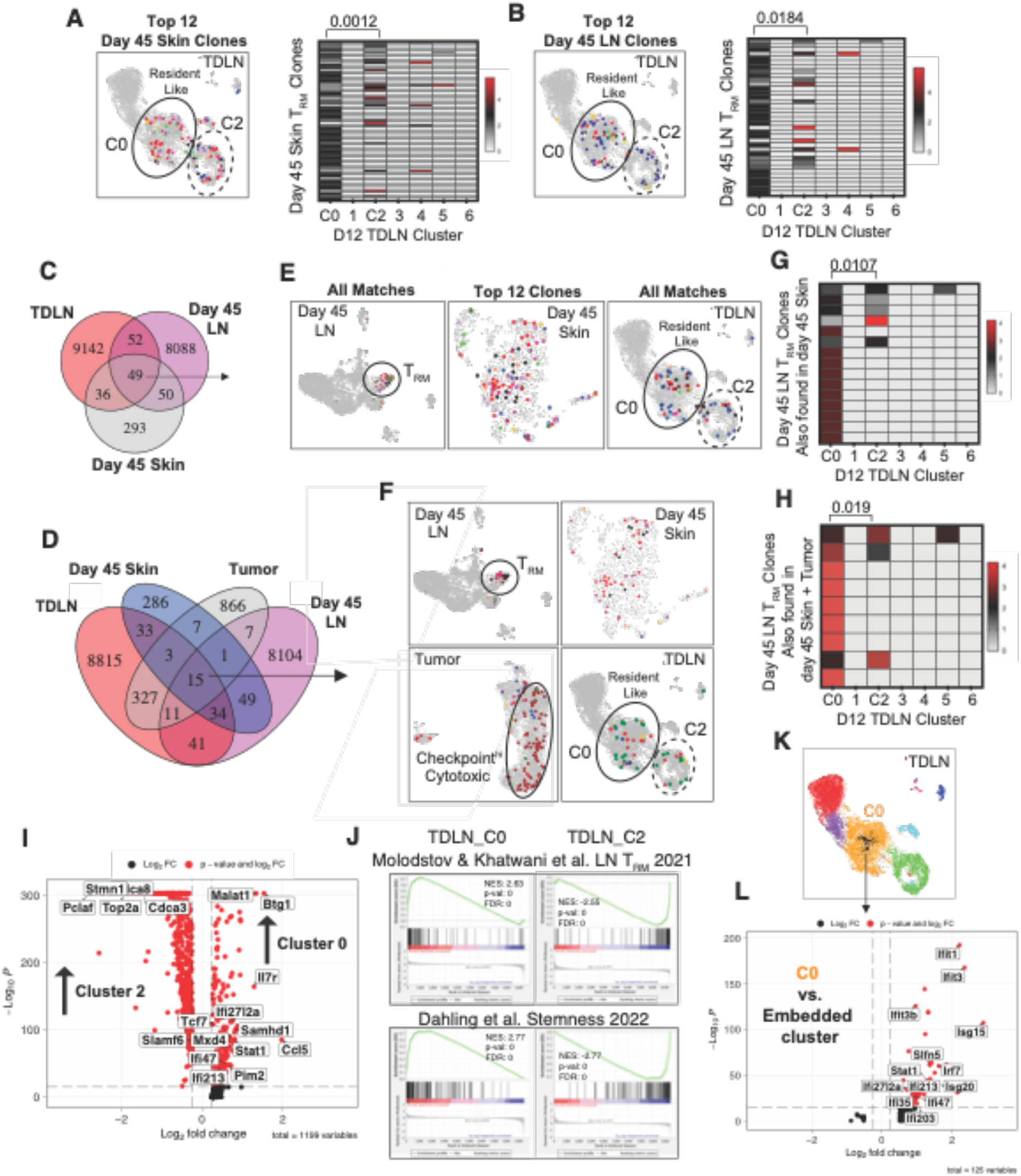
T**r**m **clonotypes originate from resident/stem-like precursors that sense type-1 IFN.** (**A-B**) (**Left**) Top 12 clonotypes matched between d12 TDLN and d45 skin (**A**), or d45 LN (**B**). (**Right**) Clonotype enrichment across the seven d12 TDLN clusters for matched Trm clonotypes found in d45 skin (A), or d45 LNs (B). (**C-D**) Venn diagrams displaying clonal matches across the indicated tissue compartments. (**E-F**) scRNA UMAP plots highlighting these matched clonotypes across their tissues of occupancy. (**G-H**) Clonotype enrichment across the seven d12 TDLN clusters for matched clonotypes that were found in d45 skin and LNs (**G**), or d45 skin and LNs, as well as in tumors (**H**). (**I**) Volcano plot of differentially expressed genes between d12 TDLN C0 and C2; log2FC >0.25. (**J**) GSEA plots showing net enrichment score (NES) of indicated gene signatures in the indicated d12 TDLN clusters. Reference dataset used is from Molodtsov and Khatwani et al. 2021 (LN Trm) and Dahling et al. 2022 (Stemness). (**K-L**) d12 TDLN scRNA UMAP and volcano plot showing differentially expressed genes between TDLN_C0 (orange) and an embedded type-1 IFN sensing cluster (black); log2FC >0.25.

We also wished to understand the features of pre-Trm clonotypes that were able to occupy multiple tissue locations. Indeed, 49 clonotypes shared between LN and skin on day 45, had counterparts in TDLNs on d12 (Fig 5C). As expected, these clonotypes mapped to all T_RM_ clusters in vitiligo skin, and almost exclusively to the T_RM_ cluster in LN. Moreover, fifteen of the clonotypes that formed Trm in skin and LNs also occupied tumors, in which they clustered based on high expression of activation, negative checkpoints, and effector transcripts (Fig. 5D, and to Fig. 3E). Projecting each of these multi-tissue distributed Trm clonotypes onto d12 TDLN clusters, we found enrichment similar to that described above, with significantly stronger occupancy in TDLN_C0 compared with the other TDLN clusters (Fig 5-H). Thus, TDLN_C0 appeared synonymous with a multipotent pre-Trm fate.

Differential gene expression analysis between TDLN_C0 and the next most represented TDLN cluster (C3) highlighted that TDLN_C0 had highest expression of genes associated with TGF-β signaling (*Btg1, Malat1, Samhd1*), and stemness (*Slamf6, Tcf7, Il7r*), and lowest expression of proliferative and cytotoxic markers (*Stmn1, Cdca8, Top2a, Cdca3, Pclaf, Gzma*) (Fig 5I). Gene set enrichment analysis (GSEA) further showed that only TDLN_C0 enriched significantly for a lymph node T_RM_ signature (*23*), and a stemness-associated gene signature (*67*) (Fig 5J). Interestingly, TDLN_C0 also contained an embedded cluster marked entirely by expression of type 1 IFN sensing genes (*Ifit1, Ifit3, Ifit3b, Isg15, Irf7, Isg20, Stat1*) (Fig 5K, L). Transcription factor (TF) enrichment and gene ontology pathway analyses revealed that TDLN cluster 0 uniquely enriched in TFs associated with stemness, tissue residency and type-1 IFN signaling (Fig. S7). Based on this, we sought to explore a role for type-1 IFN signals in enforcing the Trm fate.

### Plasmacytoid dendritic cell-derived type-1 IFN promotes Trm generation in lymph nodes

Type-1 IFN can promote memory formation by limiting proliferation and terminal differentiation of CD8 effector T cells (*68, 69*). We used Pmel cells expressing an *Mx1-Gfp* reporter gene construct (*70*) to directly assess the early time course of IFN-sensing in TDLNs. Sensing was detected as early as days 3-7 of tumor growth, reached peak levels between days 7 and 12, and then gradually declined after tumor excision on day 12 (Fig. 6A). Interestingly, GFP+ IFN-sensing Pmel cells were more likely than their GFP-non-sensing counterparts to adopt a CD103^+^CD69^+^ phenotype in TDLNs (Fig. 6B), consistent with a pre-Trm state. Plasmacytoid dendritic cells (pDCs) are major producers of type 1 interferon in viral models and in cancer (*71, 72*). Accordingly, when we treated mice to deplete pDCs (Fig. S8 A-C), we observed a significant reduction in the type-1 IFN-sensing pmel cell population in TDLNs (Fig. 6C, D). Thus pDC-derived IFN was sensed by tumor-specific T cells in TDLNs, and was associated with the acquisition of a pre-Trm state.

**Figure 6:**
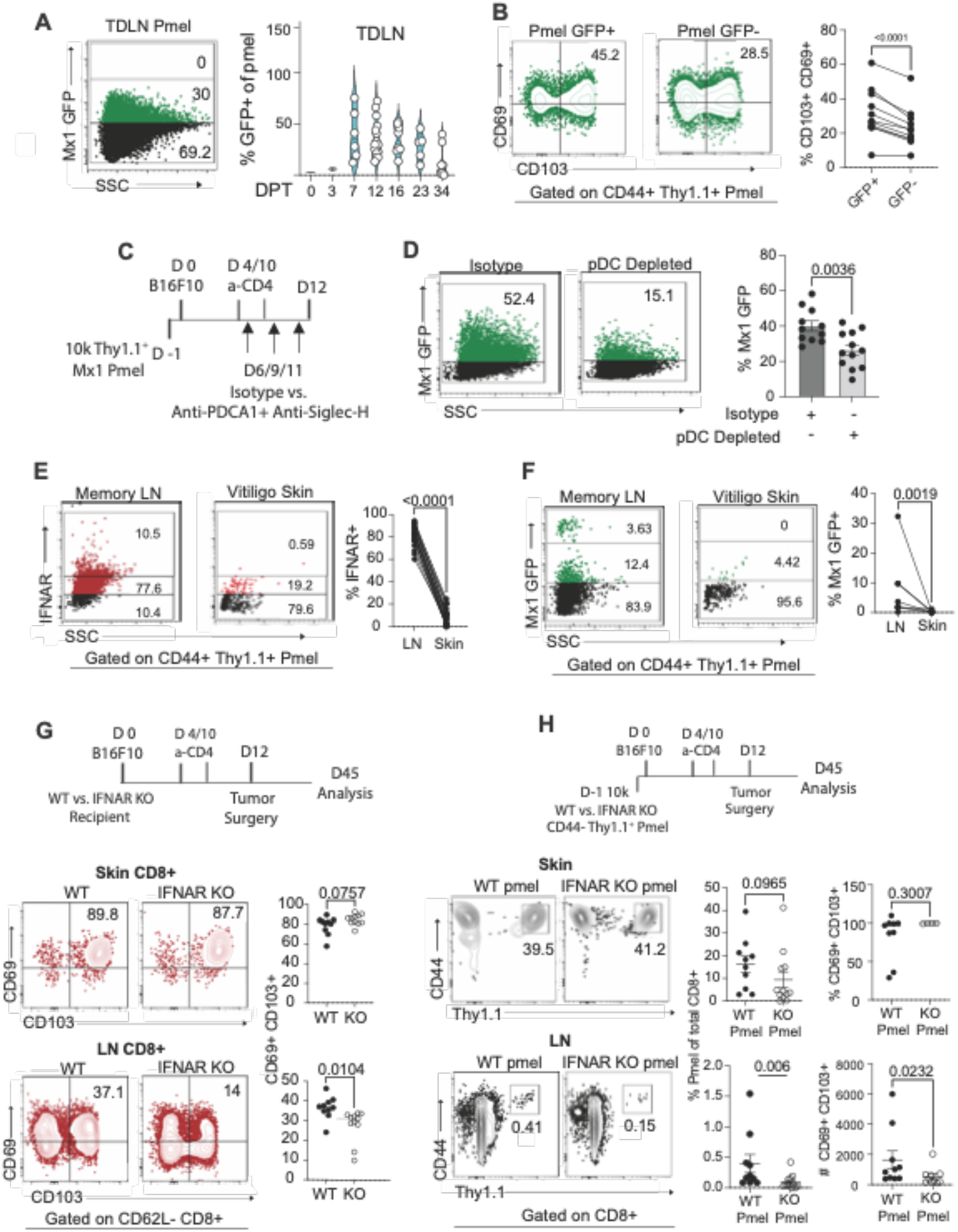
L**o**ss **of type 1 interferon sensing in tumor Ag-specific T cells impairs Trm generation in LN.** (**A**) proportion of Mx1 GFP+ gated on CD8+ CD44+ Mx1 Thy1.1+ pmel cells in TDLN at different time intervals (**B**) Proportion of CD69+/-CD103+/-GFP+ (IFN-sensing) vs. GFP-(non-sensing) pmel cells in d12 TDLN. (**C**) Schematic showing pDC depletion regimen. (**D**) proportion of Mx1 GFP+ pmel cells in d12 TDLN across treatment groups. (**E**) Proportion of IFNAR+ pmel cells in LN (left) and skin (right) 30–60 days after tumor surgery (memory). (**F**) Proportion of Mx1 GFP+ pmel cells in LN (left) and skin (right) 30–60 days after tumor excision surgery (memory) (**G, H**) Schematic showing B16 melanoma neoadjuvant treatment model in either WT or *Ifnar1-/-* mice (**G**) or in WT mice with adoptively transferred WT pmel or *Ifnar1-/-* pmel cells (**H**). (**G-Bottom**) Proportion of CD69+/– and CD103+/– total endogenous CD8 T cells, 30-60 days after tumor surgery (memory) with cumulative data (from 2 experiments) on the right. (**H-Bottom**) Proportion of Thy1.1+ pmel cells in skin and LN 30–60 days after tumor surgery, with cumulative data (from 2 experiments) on the right. (**H-right**) Proportion (skin) or absolute count (LN) of CD69+ CD103+ pmel cells. Symbols represent individual mice; horizontal lines depict means; significance was determined by unpaired (**G, H**) or paired (**B, E, F**) t test, with n.s. (non-significant) denoting p > 0.05. Flow plots depict representative data from two independent experiments; (n=5-12 mice/group); mean ± SEM.

Our prior RNAseq analysis of pmel cells had revealed a type-1 IFN sensing population of differentiated Trm cells in LN, but not in skin, on day 45 (*23*)(Fig. 1B). Accordingly, staining of interferon receptor alpha (IFNAR) revealed much higher expression on Trm cells in LNs as compared with their counterparts in the skin (Fig. 6E). Indeed, under steady state conditions, Pmel *Mx1-GFP* Trm cells in LNs also sensed higher levels of type-1 IFN than their counterparts in the skin (Fig. 6F). Thus, IFN responsiveness was indeed a unique defining feature of the Trm compartment in LNs.

Finally, we sought to determine a role for type-1 IFN in generating the Trm response. First, tumor-bearing *Ifnar-/-* mice were treated with anti-CD4 neoadjuvant therapy, and rested for 1 month, prior to analysis of the endogenous CD103^+^CD69^+^CD8^+^ Trm compartment (Fig. 6G). Interestingly, whereas no difference was observed in the Trm compartment in the skin, *Ifnar-/-* mice exhibited a significant reduction in the generation of CD103+CD69+CD8+ T cell populations in LNs (Fig. 6G). A similar reduction in LN Trm was observed in mice that received pmel cells and were treated with a blocking antibody to IFNAR (Fig. S8D-F). Of note, these reductions were not associated with changes in the severity of melanoma-associated vitiligo which was unaffected by IFNAR deficiency (Fig. S8G). Lastly, to segregate the defect in IFN sensing to tumor-antigen-specific T cells, we adoptively transferred mice with wild-type vs. *Ifnar-/-* pmel T cells and again inoculated tumors and administered neoadjuvant treatment to induce memory (Fig. 6H). Affirming our earlier findings, *Ifnar-/-* pmel proportions and their expression of CD103 and CD69 was unaltered in the skin, but proportions and absolute numbers of CD69^+^ CD103^+^ pmel Trm cells significantly reduced in LNs (Fig. 6H). These results reveal the importance of type-1 IFN/IFNAR signaling in promoting Trm generation in LNs. Taken as a whole, our findings support a model whereby pre-Trm populations that sense type-1 IFN can efficiently commit to both skin and LN Trm fates.

## DISCUSSION

Developing durable host-wide immunity against cancer is thought to be critical for successful outcomes. However, the fact that cancers metastasize necessitates the function of memory T cells in diverse tissue microenvironments. We previously showed that protective melanoma-specific T_RM_ cells can be generated in the skin (*44*), and in tumor-draining lymph nodes–– a site of early metastasis (*23*). The current study provides a new understanding of the transcriptional properties and molecular signals that promote such long-lasting T_RM_ responses to cancer. Leveraging scTCR seq for real time clonal tracing of populations enabled us to define the features of T_RM_ precursors. These cells expressed genes associated with tissue residency, but notably also acquired features of stemness and type 1 interferon sensing, the latter of which uniquely promoted Trm establishment in LNs. Thus, by linking T cell transcriptional status during primary tumor growth with protective Trm compartments in the periphery, we have defined the features of precursors that become protective Trm in the setting of cancer.

Prior studies have established that cues within tissue, such as TGF-β, IL-7 and IL-15 initiate T_RM_ fate after tissue entry (*29, 73*). However, more recent work has revealed that Trm fate is already present in circulating Teff cells (*33*). Our studies here are consistent with the later of these two models. Our finding that Trm precursors seed their tissues of occupancy early is also consistent with reports from viral infection models (*26, 30, 33*). While our trafficking studies revealed no requirement for Trm cell migration after day 12, we found that earlier trips through the tumor do occur. This is consistent with retrograde T cell migration from tumors to TDLNs reported by other investigators (*74, 75*). While prior work in human ovarian cancer reported that TCF1+ cell in the tumor microenvironment give rise to Trm populations (*76*), we did not observe clonal relationships between TCF7+ stem-like TILs in tumors and Trm cells in LNs. Thus, a tumor-derived pre-T_RM_ population seems unlikely in our model. Regardless, we found that Trm commitment was not absolute at the precursor stage. Locking precursors in tissues enhanced Trm generation, underscoring the importance of tissue-derived signals for completing the Trm differentiation program.

Early and late analyses of pmel cell populations revealed that certain transcriptional, epigenetic, and phenotypic features of Trm cells are adopted at the Teff stage, in the TDLN. However, clonal tracing of the open repertoire was required to conclusively identify this pre-Trm state. This approach provided resolution to observe polyclonal T cell populations and classify them based on their propensity to form Trm populations. In accordance with recent studies (*33*), we show that pre-Trm cells are high for residency and stemness markers, as well as markers associated with TGF-b signaling (*63*). Interestingly, we found no evidence of *Hobit* expression, despite its implication in viral-specific Trm (*39, 42*). Instead, we found *Id2* and *Id3*–– transcription known to promote T_RM_ differentiation (*77, 78*) to be abundantly expressed pre-Trm pmel cells.

Relative to other memory fates, the Trm fate appeared dominant in this neoadjuvant treatment model, with clonotypes that differentiated into Trm cells showing little evidence of taking on other fates. Progenitor/exhausted-like cells in LNs were clonally expanded, but they derived from a distinct clonal repertoire. That said, pre-Trm clonotypes had progeny that simultaneously formed Trm in multiple locations, highlighting the existence of multipotent Trm precursors. Our data suggest a model of early fate commitment to Trm at the clonal level, with subsequent differentiation to Trm, but with molecular signals—such as type-1 IFN— fine-tuning residence in specific locations. One remaining question is how TCR affinity may affect such fate decisions. Recent studies argue for both positive (*79, 80*), and negative (*80, 81*), association between TCR signal strength and Trm generation. Our model of immunity to melanoma with associated vitiligo is dominated by recognition of tumor/self-antigens, which are mainly of low affinity, although it is not known whether this naturally skews cells to a Trm fate.

We also found that clonotypes with confirmed propensity to become Trm failed to do so in the TME. While aggressive tumor growth may not provide enough time for Trm differentiation, one would expect that some Trm features would have been apparent, particularly as B16 tumors produce high levels of TGF-b. Recent studies also report the absence of a *bona fide* Trm phenotype in mouse tumors, although T cells with alternate phenotypes still stably resided in tumors (*82*). It will be important to understand what factors promote Trm generation in the TME, as Trm populations that clonally and transcriptionally overlap with vitiligo-affected skin, are in fact present in human melanoma tumors (*83*).

Whereas type-1 IFN signals have not previously been implicated in promoting the Trm fate, IFN is known to drive T cell survival and memory formation in the setting of viral infection (*68, 69, 84*). Prior work shows that Type-1 IFN signals results from STING activation in TDLNs (*85*). Moreover, IFN is known to provide a “stay” signal for Teff cells in LNs by upregulating their expression of CD69 and antagonizing S1PR1 (*86*). As we find that differentiated Trm cells continue to express IFNAR and sense IFN within LNs, it remains unclear whether type-1 IFN signals are sufficient during programming, or if ongoing sensing is required. Of note, a robust type 1 IFN signature has been reported as an early event in vitiligo pathogenesis (*87*). While we identified pDCs as a source of type-1 IFN, type-1 DCs are also major producers in the setting of cancer (*88*). Specialized niches within the TDLN could potentially facilitate provision of type-1 IFN to a subset of Teff cells that then adopt a multipotent Trm fate.

In summary, this research establishes how Trm fate decisions are made, over time and across tissues, during an anti-tumor response. Prior to this, traits that define multipotent Trm precursors remained unclear. Knowledge of such fundamental properties may be critical for future advances in adoptive cell therapy, particularly as CAR T cells skewed to the Trm fate with TGF-b, were recently shown to have enhanced anti-tumor efficacy (*89*). The provision of additional stemness-promoting factors and molecular signals such as type 1-IFN may help to generate more broadly distributed and resilient Trm populations.

## MATERIALS AND METHODS

### Experimental model

Experiments on living animals. Mice of the strain C57BL/6 were acquired from Charles River Laboratories between the ages of eight and ten weeks and used in the tests between the ages of eight and twelve weeks. Ten-week-old Pmel-1 (pmel) mice were produced in-house (originally a gift from N. Restifo; The National Cancer Institute) on a Thy1.1 congenic background and used for experiments. In-house breeding of Kaede-GFP and Mx1-GFP reporter mice, *Ifnar1* −/− mice (see table S1) was accomplished by the acquisition of these strains from the Jackson Laboratory. Dartmouth College’s Comparative Medicine and Research facility ensured that all animals were maintained in a pathogen-free environment. Mice were kept in a controlled temperature environment with regular light and dark cycles in isolator caging units. All research with animals at Dartmouth was conducted in accordance with the recommendations of the university’s animal care and use committee. Each experiment’s sample size and primary outcome were predetermined by a systematic review of the literature on detecting analogous biological effects. Outliers, along with the rest of the analyzed samples, were incorporated into the results. The picture legends and the text that follows provide more specifics about the experimental setup and data processing.

### Cell lines and antibodies

To ensure consistent growth in the dermis, the B16-F10 mouse melanoma cell line was passed intradermally (i.d.) fifteen times in C57BL/6 mice after being obtained from Isaiah Fidler at the MD Anderson Cancer Center (*44*). All intradermal primary tumors were derived from this line. IDEXX BioAnalytics confirmed that B16 lines to be free of viruses and mycobacterium (Columbia, MO). Bioreactor supernatants from hybridoma cell lines (American Type Culture Collection) were used to produce depleting anti-CD4 (mAb clone GK1.5), which was then injected IP at doses of 200 μg. unless specified otherwise, all fluorescent antibodies used for flow cytometry were provided by from Biolegend or eBiosciences.

### Neoadjuvant treatment of B16 melanoma with anti-CD4

As we described previously (*90*), C57BL/6 mice, and various strain on the C57BL/6 background, were injected intradermally on the right flank with 1.75 × 105 B16 cells on day 0 and treated with anti-CD4 mAb intravenously on days 4 and 10 to efficiently deplete regulatory (Treg) cells. On day 12, dermal tumors were surgically excised under isofluorane anesthesia. Mice with recurrent primary tumors after surgery (5%) were removed from the study. Study animals that developed vitiligo (∼70-80%, as previously demonstrated (*91*)) were used for analysis at memory timepoints. As previously described, mice with both localized (at the surgical site) and disseminated (beyond the surgical site) vitiligo were used interchangeably in experiments (*44, 91*).

### Pmel cell adoptive transfer

Naïve CD8+Thy1.1+ pmel cells were isolated from 8–10 week old pmel T cell receptor transgenic mice. CD44-cells were purified using Mouse Naive CD8 T cell isolation kit from Stemcell. Cell purity and naive phenotype was confirmed by flow cytometry before injecting them into recipients. Unless otherwise stated, 10,000 naive pmel cells were injected retro-orbitally one day prior to the tumor transplantation.

### FTY720 treatment

Mice received daily i.p. injections of FTY720 (2-amino-2-(2[4-octylphenyl]ethyl)-1,3-propanediol hydrochloride, Cayman Chemical), (1 mg/kg) or through drinking water that was replenished every 2 days until the study endpoint.

### TCR clonal tracing in individual mice

Melanoma associated vitiligo was induced in mice without adoptive transfer of sentinel Pmel T cells in order to monitor the endogenous CD8+ T cell repertoire. On day 12 after tumor transplantation on ipsilateral and contralateral flanks, tumor surgery was performed on both flanks and at the same time, both TDLN (ipsilateral TDLN and contralateral TDLN) were excised. TDLNs and tumors (priming) were processed and stained with a combination or hashtag and surface antibodies and CD8+ CD44+ T cells were sorted and submitted for paired scRNA/TCR sequencing. Mice that underwent this surgery were monitored for vitiligo establishment and only mice that developed autoimmune vitiligo were considered for TCR fate mapping studies at memory time point. From these mice, distal axillary/brachial lymph nodes and matched vitiligo skin were processed for cell sorting. CD8+ CD44+ CD62L^low/int^ T cells were sorted from LN and CD8+ CD44+ T cells were sorted from vitiligo skin and submitted for paired scRNA/TCR sequencing. In another similar experiment, ipsilateral LN and both tumors were excised on the day of tumor surgery (d12 priming) and mice were allowed to develop vitiligo. Only vitiligo-stricken mice were used at memory and contralateral LN and matched vitiligo skin were processed for cell sorting. Tissues were processed as described above and stained with a combination or hashtag and surface antibodies. CD8+ CD44+ T cells were sorted and submitted for paired scRNA/TCR sequencing at both priming and memory time points.

### Photoconversion studies with KAEDE-pmel cells

*In situ* KAEDE-pmel cell photoconversion in tumors was accomplished with a 405 nm UV light source by exposing the tissue of interest to the light source for 5-10 minutes at 50% intensity. For photoconverting skin after tumor excision, tissue was exposed to UV light for 15 minutes while the probe moved continuously to maximize photoconversion over a large area of skin. For the duration of the exposure, the area adjacent to the tissue of interest was covered with opaque foil to prevent the light source from dispersing. In the case of tumor photoconversion, for example, the area surrounding the tumor was covered with foil, leaving only the tumor exposed to UV radiation. This procedure was repeated for each photoconversion round. No tissue hypertrophy or wounding was observed.

### Mouse tissue harvesting and digest

To ensure memory formation, all mice were sacrificed at least 28 days after primary tumor excision. Spleens and lymph nodes were mechanically dissociated. Unless otherwise specified, analyses were performed on a section of depigmented epidermis from the right flank (the primary site of tumor excision). A 2 cm^2^ patch of skin was minced and incubated in 3 mg/mL collagenase Type IV (Worthington Biochemical Corporation) and 2 mg/mL DNase (Sigma-Aldrich) in HBSS at 37 °C for 45 minutes with stirring. Tumors were processed using a percoll gradient, and the leukocyte population/layer was collected at the interface between 40% and 80% discontinuous gradient. Single cell suspensions from various tissues were processed for flow cytometry or cell sorting.

### Analysis of T cells by flow cytometry

Live-dead staining was performed for 15 minutes in cold PBS prior to antibody staining. After thorough washing, anti-mouse FC blocking Ab (Bioxcell) was applied on ice for 15 minutes to samples prepared as described above. Antibodies (see table S2) were used in different combinations to stain cells for surface markers. After 30 minutes of incubation at 4°C, cells were washed twice with Flow buffer (1X PBS containing 0.2% BSA) and fixed overnight with 1-2% formaldehyde. Flow cytometry was performed on a Bio-Rad ZE5 Cell Analyzer, and data was analyzed with Flowjo v10 software (Treestar Inc). Pmel cell phenotypes are only reported for confined populations with at least 40 events.

### Fluorescence Activated Cell Sorting (FACS) of T cells for scRNA/scTCR seq analyses

Tissues were digested as described above, with tissue specimens pooled from n>=5 mice. Single cell suspensions were prepared for each tissue and stained with anti-CD8 (53–6.7), anti-Thy1.1 (OX-7), BV510-conjugated anti-mouse CD62L (MEL-14) and anti-CD44 (IM7). In mice that did not receive pmel cells, endogenous CD8+CD44hi or CD8+ CD44+ CD62l^int/low^ T cells were sorted to >90% purity using an SH800 cell sorter (SONY). For single cell (sc) RNA-seq/ATAC-seq experiments, CD8+Thy1.1+ pmel cells were isolated from tumor bearing mice on day 12 post tumor transplantation or mice with vitiligo 45 days following primary tumor excision.

### Single-cell RNAseq, TCRseq and ATACseq

For each sample, cells were FACS sorted into a single tube to minimize sample loss. In order to obtain the maximum number of cells from limited material, the entire sorted volume (up to 33.8ul) was loaded onto a single lane of a Single Cell A chip and processed on a Chromium instrument (10X Genomics). Libraries were prepared using the 10X Single cell 3’ V2 chemistry according to the manufacturer’s protocol. Libraries underwent quality control by Fragment Analyzer and Qubit (ThermoFisher) to determine size distribution and the quantity of the libraries prepared. Amplified cDNA and TCR-specific libraries were prepared following the standard 10× procedure to generate libraries for Illumina sequencing. Samples were uniquely barcoded, pooled and sequenced across multiple Illumina NextSeq500High Output runs to generate 50,000 and 5000 reads/cell for gene expression and TCR libraries, respectively. Paired end sequencing was performed using 26 cycles for read 1 to decode the 16bp cell barcode and 10bp UMI sequences, and 98 cycles for read 2 corresponding to the transcript sequence. Raw sequencing data was processed through the Cell Ranger v3.0 pipeline (10× Genomics) using the mouse reference genome mm10 to generate gene expression matrices for single cell 5’ RNA-seq data and, in some instances, fully reconstructed, paired TRA/TRB sequences.

For scATAC-seq, nuclei were isolated from single cell suspensions following the 10x Genomics protocol (CG000169) and loaded onto a Chromium NextGEM Chip H for capture. Libraries were prepared according to the 10x Genomics Single Cell ATAC v1.1 protocol (CG000209) and pooled for sequencing on a NextSeq500 instrument (Illumina) (Read1: 50bp; Read2: 50bp; Index1: 8bp; Index2: 16bp). FASTQ files were processed using cellranger v1.2.0.

### Single cell RNAseq, ATACseq, and TCRseq data analysis

As previously published (*92, 93*), the “Seurat v3” R package was applied to filter out low-quality cells, normalize gene expression profiles, and cluster cells. Cells expressing >5% mitochondrial gene counts or expressing less than 500 genes were discarded using the FilterCells function. Then, the NormalizeData function was applied to normalize, and log transformed the raw counts for each cell based on the library size. Clustering of cells was performed using Seurat v3.0 pipeline and resolution was set based on differentially expressed genes. Data involving more than one tissue or time point sequenced on different lanes was analyzed after merging the datasets followed by batch correction using the Seurat “vars.to.regress” function. For single cell ATAC sequencing analysis, Signac and Seurat pipeline were applied for performing QC metrics, including nucleosomal signal and TSS enrichment score analysis. This was followed by normalization for nonlinear reduction and clustering. Gene activity matrices were created for each sample, and chromatin accessibility peaks were identified for genes for plotting genomic regions. For scTCR analysis, the 10× Cellranger VdJ pipeline was used to determine each TCR α-chain and β-chain CDR3 sequence for a corresponding cell. Only productive TCRs were used to identify TCR clonotypes. Identical CDR3 nucleotide sequences were required for cells to be identified as a matched clonotype. The scRepertoire pipeline was used to calculate clonal overlay and occupancy (*94*).

### Gene set enrichment analysis (GSEA) and Transcriptional Factor Analysis

Published gene signatures of interest, (*23, 67*) were utilized to run GSEA analysis on clusters and tissues of interest. These gene signatures were added to the ‘c2.all.v6.2.symbols’ gene sets collection from the MSigDB database for GSEA analysis. The GSEA 4.0.3 software was used to conduct pre-ranked GSEA. Using the FindAllMarkers function in Seurat with the parameters logfc. thereshold=0.01 and min.pct=100%, the genes of each cluster utilized for the analysis were identified. The log2FC value was utilized as the metric for ranking. For transcription factor analysis, differentially expressed genes for indicated tissues were used as gene input list in ChEA3 pipeline (ref) and mean rank was used to identify top TFs enriched in cluster of interest.

### Enrichment score analysis

To determine if a given group of TCR cell clonotypes was enriched in a particular UMAP cluster, a ‘Enrichment score’ was calculated using a hypergeometric distribution, while removing potential biases resulting from cell number differences between mice. For instance, from a given tissue ‘T’, a total of ‘N’ cells was sequenced, and for a given T cell clone, ‘x’ cells belonged to this clonotype. Among these cells, ‘y’ total cells were in cluster A, which contained ‘m’ number of cells from ‘T’. Enrichment score for clonotype in cluster A is then computed as = y x/N × m

### Statistical analyses

The GraphPad Prism 5 software was utilized for statistical analysis (GraphPad Software Inc). The Gaussian distribution of the data was examined using the D’Agostino & Pearson test and the Shapiro-Wilk test. When normally distributed according to one of these tests, statistical differences were analyzed using the unpaired t-test to compare two groups from different mice and the paired t-test to compare two values from the same mouse. When data was not normally distributed or sample size was insufficient to determine the distribution, the Mann-Whitney test was used to compare two unrelated groups, whereas the Wilcoxon matched pairs test was used to compare two values from the same mouse. t tests were two-sided, and a p value 0.05 was regarded as statistically significant.

### Supplementary Materials

Tables S1 and S2

Figs. S1-S8

## Funding

National Institutes of Health grants R01CA22502 to MJT, R01CA254042 to MJT and YHH, P20GM130454 to FWK, 5P30CA023108 to MJT and FWK, R01CA269455 to PCR, and T32 AI00763 to DER.

O. Ross McIntyre, M.D. Endowment to MJT.

Knights of the York Cross of Honour philanthropic support.

## Author contributions

Conceptualization: NK, MJT

Methodology: NK, FWK, MJT

Investigation: NK, TGS, JH, CM, NM, AKM, DER, AH, OW, FWK

Formal Analysis: NK, MJT

Bioinformatics Analysis: NK, TGS, JH, OW, FWK Supervision: MJT

Writing – original draft: NK, MJT

Writing – review & editing: NK, YHH, PCR, FWK, MJT

## Data and materials availability

Single cell transcriptomics data, including scRNA, scTCR and scATAC have been uploaded to the Gene Expression Omnibus (GEO) under accession number GSE236285. All data needed to evaluate the conclusions in the paper are present in the paper or the supplementary materials.

## REFERENCES

1. H. Dillekas, M. S. Rogers, O. Straume, Are 90% of deaths from cancer caused by metastases? Cancer Med 8, 5574–5576 (2019).

2. D. Hanahan, R. A. Weinberg, Hallmarks of cancer: the next generation. Cell 144, 646–674 (2011).

3. V. Shankaran et al., IFNgamma and lymphocytes prevent primary tumour development and shape tumour immunogenicity. Nature 410, 1107–1111 (2001).

4. K. J. Hiam-Galvez, B. M. Allen, M. H. Spitzer, Systemic immunity in cancer. Nat Rev Cancer 21, 345–359 (2021).

5. N. P. Restifo, M. E. Dudley, S. A. Rosenberg, Adoptive immunotherapy for cancer: harnessing the T cell response. Nat Rev Immunol 12, 269–281 (2012).

6. C. L. Ventola, Cancer Immunotherapy, Part 1: Current Strategies and Agents. P T 42, 375–383 (2017).

7. J. Han, N. Khatwani, T. G. Searles, M. J. Turk, C. V. Angeles, Memory CD8(+) T cell responses to cancer. Semin Immunol 49, 101435 (2020).

8. D. S. Chen, I. Mellman, Elements of cancer immunity and the cancer-immune set point. Nature 541, 321–330 (2017).

9. S. L. Park, T. Gebhardt, L. K. Mackay, Tissue-Resident Memory T Cells in Cancer Immunosurveillance. Trends Immunol 40, 735–747 (2019).

10. R. Ahmed, D. Gray, Immunological memory and protective immunity: understanding their relation. Science 272, 54–60 (1996).

11. S. M. Kaech, W. Cui, Transcriptional control of effector and memory CD8+ T cell differentiation. Nat Rev Immunol 12, 749–761 (2012).

12. S. N. Mueller, T. Gebhardt, F. R. Carbone, W. R. Heath, Memory T cell subsets, migration patterns, and tissue residence. Annu Rev Immunol 31, 137–161 (2013).

13. S. N. Mueller, L. K. Mackay, Tissue-resident memory T cells: local specialists in immune defence. Nat Rev Immunol 16, 79–89 (2016).

14. A. Molodtsov, M. J. Turk, Tissue Resident CD8 Memory T Cell Responses in Cancer and Autoimmunity. Front Immunol 9, 2810 (2018).

15. J. M. Schenkel, K. A. Fraser, V. Vezys, D. Masopust, Sensing and alarm function of resident memory CD8(+) T cells. Nat Immunol 14, 509–513 (2013).

16. J. M. Schenkel et al., T cell memory. Resident memory CD8 T cells trigger protective innate and adaptive immune responses. Science 346, 98–101 (2014).

17. T. Wang, Y. Shen, S. Luyten, Y. Yang, X. Jiang, Tissue-resident memory CD8(+) T cells in cancer immunology and immunotherapy. Pharmacol Res 159, 104876 (2020).

18. T. Tsuji, J. Matsuzaki, K. Odunsi, Tissue residency of memory CD8(+) T cells matters in shaping immunogenicity of ovarian cancer. Cancer Cell 40, 452–454 (2022).

19. S. L. Park et al., Tissue-resident memory CD8(+) T cells promote melanoma-immune equilibrium in skin. Nature 565, 366–371 (2019).

20. X. Jiang et al., Skin infection generates non-migratory memory CD8+ T(RM) cells providing global skin immunity. Nature 483, 227–231 (2012).

21. T. Gebhardt et al., Memory T cells in nonlymphoid tissue that provide enhanced local immunity during infection with herpes simplex virus. Nat Immunol 10, 524–530 (2009).

22. M. Enamorado et al., Enhanced anti-tumour immunity requires the interplay between resident and circulating memory CD8(+) T cells. Nat Commun 8, 16073 (2017).

23. A. K. Molodtsov et al., Resident memory CD8(+) T cells in regional lymph nodes mediate immunity to metastatic melanoma. Immunity 54, 2117–2132 e2117 (2021).

24. F. Mami-Chouaib et al., Resident memory T cells, critical components in tumor immunology. J Immunother Cancer 6, 87 (2018).

25. D. J. Craig et al., Resident Memory T Cells and Their Effect on Cancer. Vaccines (Basel) 8, (2020).

26. F. E. Dijkgraaf, L. Kok, T. N. M. Schumacher, Formation of Tissue-Resident CD8(+) T-Cell Memory. Cold Spring Harb Perspect Biol 13, (2021).

27. J. J. Milner et al., Runx3 programs CD8(+) T cell residency in non-lymphoid tissues and tumours. Nature 552, 253–257 (2017).

28. J. J. Milner, A. W. Goldrath, Transcriptional programming of tissue-resident memory CD8(+) T cells. Curr Opin Immunol 51, 162–169 (2018).

29. L. K. Mackay et al., The developmental pathway for CD103(+)CD8+ tissue-resident memory T cells of skin. Nat Immunol 14, 1294–1301 (2013).

30. N. S. Kurd, et al., Early precursors and molecular determinants of tissue-resident memory CD8(+) T lymphocytes revealed by single-cell RNA sequencing. Sci Immunol 5, (2020).

31. L. Kok, D. Masopust, T. N. Schumacher, The precursors of CD8(+) tissue resident memory T cells: from lymphoid organs to infected tissues. Nat Rev Immunol 22, 283–293 (2022).

32. B. S. Sheridan et al., Oral infection drives a distinct population of intestinal resident memory CD8(+) T cells with enhanced protective function. Immunity 40, 747–757 (2014).

33. L. Kok et al., A committed tissue-resident memory T cell precursor within the circulating CD8+ effector T cell pool. J Exp Med 217, (2020).

34. V. Mani et al., Migratory DCs activate TGF-beta to precondition naive CD8(+) T cells for tissue-resident memory fate. Science 366, (2019).

35. O. Gaide et al., Common clonal origin of central and resident memory T cells following skin immunization. Nat Med 21, 647–653 (2015).

36. C. I. Yu et al., Human CD1c+ dendritic cells drive the differentiation of CD103+ CD8+ mucosal effector T cells via the cytokine TGF-beta. Immunity 38, 818–830 (2013).

37. S. Iborra et al., Optimal Generation of Tissue-Resident but Not Circulating Memory T Cells during Viral Infection Requires Crosspriming by DNGR-1(+) Dendritic Cells. Immunity 45, 847–860 (2016).

38. P. Bourdely et al., Transcriptional and Functional Analysis of CD1c(+) Human Dendritic Cells Identifies a CD163(+) Subset Priming CD8(+)CD103(+) T Cells. Immunity 53, 335–352 e338 (2020).

39. L. Parga-Vidal, et al., Hobit identifies tissue-resident memory T cell precursors that are regulated by Eomes. Sci Immunol 6, (2021).

40. R. T. Sowell, M. Rogozinska, C. E. Nelson, V. Vezys, A. L. Marzo, Cutting edge: generation of effector cells that localize to mucosal tissues and form resident memory CD8 T cells is controlled by mTOR. J Immunol 193, 2067–2071 (2014).

41. N. A. Kragten et al., Hobit and Blimp-1 instruct the differentiation of iNKT cells into resident-phenotype lymphocytes after lineage commitment. Eur J Immunol 52, 389–403 (2022).

42. L. Parga-Vidal et al., Hobit and Blimp-1 regulate T(RM) abundance after LCMV infection by suppressing tissue exit pathways of T(RM) precursors. Eur J Immunol 52, 1095–1111 (2022).

43. M. Evrard et al., Sphingosine 1-phosphate receptor 5 (S1PR5) regulates the peripheral retention of tissue-resident lymphocytes. J Exp Med 219, (2022).

44. B. T. Malik, et al., Resident memory T cells in the skin mediate durable immunity to melanoma. Sci Immunol 2, (2017).

45. A. L. Cote, K. T. Byrne, S. M. Steinberg, P. Zhang, M. J. Turk, Protective CD8 memory T cell responses to mouse melanoma are generated in the absence of CD4 T cell help. PLoS One 6, e26491 (2011).

46. C. Peng et al., Engagement of the costimulatory molecule ICOS in tissues promotes establishment of CD8(+) tissue-resident memory T cells. Immunity 55, 98–114 e115 (2022).

47. C. Alanio et al., CXCR3/CXCL10 Axis Shapes Tissue Distribution of Memory Phenotype CD8(+) T Cells in Nonimmunized Mice. J Immunol 200, 139–146 (2018).

48. M. Delacher et al., Precursors for Nonlymphoid-Tissue Treg Cells Reside in Secondary Lymphoid Organs and Are Programmed by the Transcription Factor BATF. Immunity 52, 295–312 e211 (2020).

49. J. M. Stolley et al., Retrograde migration supplies resident memory T cells to lung-draining LN after influenza infection. J Exp Med 217, (2020).

50. L. K. Beura et al., T Cells in Nonlymphoid Tissues Give Rise to Lymph-Node-Resident Memory T Cells. Immunity 48, 327–338 e325 (2018).

51. M. Tomura et al., Monitoring cellular movement in vivo with photoconvertible fluorescence protein “Kaede” transgenic mice. Proc Natl Acad Sci U S A 105, 10871–10876 (2008).

52. P. Zhang, A. L. Cote, V. C. de Vries, E. J. Usherwood, M. J. Turk, Induction of postsurgical tumor immunity and T-cell memory by a poorly immunogenic tumor. Cancer Res 67, 6468–6476 (2007).

53. V. Brinkmann et al., Fingolimod (FTY720): discovery and development of an oral drug to treat multiple sclerosis. Nat Rev Drug Discov 9, 883–897 (2010).

54. B. J. Laidlaw, E. E. Gray, Y. Zhang, F. Ramirez-Valle, J. G. Cyster, Sphingosine-1-phosphate receptor 2 restrains egress of gammadelta T cells from the skin. J Exp Med 216, 1487–1496 (2019).

55. A. N. Wein et al., CXCR6 regulates localization of tissue-resident memory CD8 T cells to the airways. J Exp Med 216, 2748–2762 (2019).

56. B. V. Kumar et al., Human Tissue-Resident Memory T Cells Are Defined by Core Transcriptional and Functional Signatures in Lymphoid and Mucosal Sites. Cell Rep 20, 2921–2934 (2017).

57. M. Nizard et al., Induction of resident memory T cells enhances the efficacy of cancer vaccine. Nat Commun 8, 15221 (2017).

58. L. M. Wakim et al., The molecular signature of tissue resident memory CD8 T cells isolated from the brain. J Immunol 189, 3462–3471 (2012).

59. R. Muthuswamy et al., CXCR6 by increasing retention of memory CD8(+) T cells in the ovarian tumor microenvironment promotes immunosurveillance and control of ovarian cancer. J Immunother Cancer 9, (2021).

60. S. F. Cai et al., Differential expression of granzyme B and C in murine cytotoxic lymphocytes. J Immunol 182, 6287–6297 (2009).

61. D. C. Yanez, S. Ross, T. Crompton, The IFITM protein family in adaptive immunity. Immunology 159, 365–372 (2020).

62. J. G. van den Boorn et al., Autoimmune destruction of skin melanocytes by perilesional T cells from vitiligo patients. J Invest Dermatol 129, 2220–2232 (2009).

63. Q. Liu et al., TGF-beta1-Induced Upregulation of MALAT1 Promotes Kazakh’s Esophageal Squamous Cell Carcinoma Invasion by EMT. J Cancer 11, 6892–6901 (2020).

64. P. A. Szabo et al., Single-cell transcriptomics of human T cells reveals tissue and activation signatures in health and disease. Nat Commun 10, 4706 (2019).

65. B. Bengsch et al., Epigenomic-Guided Mass Cytometry Profiling Reveals Disease-Specific Features of Exhausted CD8 T Cells. Immunity 48, 1029–1045 e1025 (2018).

66. M. A. Paley et al., Progenitor and terminal subsets of CD8+ T cells cooperate to contain chronic viral infection. Science 338, 1220–1225 (2012).

67. S. Dahling et al., Type 1 conventional dendritic cells maintain and guide the differentiation of precursors of exhausted T cells in distinct cellular niches. Immunity 55, 656–670 e658 (2022).

68. J. P. Huber, J. D. Farrar, Regulation of effector and memory T-cell functions by type I interferon. Immunology 132, 466–474 (2011).

69. H. J. Ramos et al., Reciprocal responsiveness to interleukin-12 and interferon-alpha specifies human CD8+ effector versus central memory T-cell fates. Blood 113, 5516–5525 (2009).

70. M. B. Uccellini, A. Garcia-Sastre, ISRE-Reporter Mouse Reveals High Basal and Induced Type I IFN Responses in Inflammatory Monocytes. Cell Rep 25, 2784–2796 e2783 (2018).

71. P. Fitzgerald-Bocarsly, J. Dai, S. Singh, Plasmacytoid dendritic cells and type I IFN: 50 years of convergent history. Cytokine Growth Factor Rev 19, 3–19 (2008).

72. B. Zhou, T. Lawrence, Y. Liang, The Role of Plasmacytoid Dendritic Cells in Cancers. Front Immunol 12, 749190 (2021).

73. T. Adachi et al., Hair follicle-derived IL-7 and IL-15 mediate skin-resident memory T cell homeostasis and lymphoma. Nat Med 21, 1272–1279 (2015).

74. M. M. Steele et al., T cell egress via lymphatic vessels is tuned by antigen encounter and limits tumor control. Nat Immunol 24, 664–675 (2023).

75. Z. Li et al., In vivo labeling reveals continuous trafficking of TCF-1+ T cells between tumor and lymphoid tissue. J Exp Med 219, (2022).

76. C. M. Anadon et al., Ovarian cancer immunogenicity is governed by a narrow subset of progenitor tissue-resident memory T cells. Cancer Cell 40, 545–557 e513 (2022).

77. J. J. Milner et al., Heterogenous Populations of Tissue-Resident CD8(+) T Cells Are Generated in Response to Infection and Malignancy. Immunity 52, 808–824 e807 (2020).

78. C. Y. Yang et al., The transcriptional regulators Id2 and Id3 control the formation of distinct memory CD8+ T cell subsets. Nat Immunol 12, 1221–1229 (2011).

79. E. L. Frost, A. E. Kersh, B. D. Evavold, A. E. Lukacher, Cutting Edge: Resident Memory CD8 T Cells Express High-Affinity TCRs. J Immunol 195, 3520–3524 (2015).

80. J. K. Fiege et al., The Impact of TCR Signal Strength on Resident Memory T Cell Formation during Influenza Virus Infection. J Immunol 203, 936–945 (2019).

81. S. Maru, G. Jin, T. D. Schell, A. E. Lukacher, TCR stimulation strength is inversely associated with establishment of functional brain-resident memory CD8 T cells during persistent viral infection. PLoS Pathog 13, e1006318 (2017).

82. N. V. Gavil, et al., Chronic antigen in solid tumors drives a distinct program of T cell residence. Sci Immunol 8, eadd5976 (2023).

83. J. Han et al., Resident and circulating memory T cells persist for years in melanoma patients with durable responses to immunotherapy. Nat Cancer 2, 300–311 (2021).

84. G. A. Kolumam, S. Thomas, L. J. Thompson, J. Sprent, K. Murali-Krishna, Type I interferons act directly on CD8 T cells to allow clonal expansion and memory formation in response to viral infection. J Exp Med 202, 637–650 (2005).

85. S. R. Woo et al., STING-dependent cytosolic DNA sensing mediates innate immune recognition of immunogenic tumors. Immunity 41, 830–842 (2014).

86. L. R. Shiow et al., CD69 acts downstream of interferon-alpha/beta to inhibit S1P1 and lymphocyte egress from lymphoid organs. Nature 440, 540–544 (2006).

87. A. Bertolotti et al., Type I interferon signature in the initiation of the immune response in vitiligo. Pigment Cell Melanoma Res 27, 398–407 (2014).

88. M. B. Fuertes et al., Host type I IFN signals are required for antitumor CD8+ T cell responses through CD8alpha+ dendritic cells. J Exp Med 208, 2005–2016 (2011).

89. I. Y. Jung et al., Tissue-resident memory CAR T cells with stem-like characteristics display enhanced efficacy against solid and liquid tumors. Cell Rep Med 4, 101053 (2023).

90. M. J. Turk et al., Concomitant tumor immunity to a poorly immunogenic melanoma is prevented by regulatory T cells. J Exp Med 200, 771–782 (2004).

91. K. T. Byrne et al., Autoimmune melanocyte destruction is required for robust CD8+ memory T cell responses to mouse melanoma. J Clin Invest 121, 1797–1809 (2011).

92. T. Stuart et al., Comprehensive Integration of Single-Cell Data. Cell 177, 1888–1902 e1821 (2019).

93. A. Butler, P. Hoffman, P. Smibert, E. Papalexi, R. Satija, Integrating single-cell transcriptomic data across different conditions, technologies, and species. Nat Biotechnol 36, 411–420 (2018).

94. N. Borcherding, N. L. Bormann, G. Kraus, scRepertoire: An R-based toolkit for single-cell immune receptor analysis. F1000Res 9, 47 (2020).

