## Supplemental Figures for "Resident memory T cell precursors in tumor draining lymph nodes require type-1 IFN for optimal differentiation": FINAL SUPPL_Khatwani et al.pdf

### SUPPLEMENTARY MATERIALS

**Table S1: Experimental models, software, and algorithms**

| | | |
|---|---|---|
| **Experimental Models: Cell Lines** | | |
| B16-F10 | Isaiah Fidler, MD Anderson Cancer Center | N/a |
| **Experimental Models: Organisms/Strains** | | |
| C57BL/6NCrl | Charles River Laboratories | Cat# CRL27, RRID: IMSR_CRL:27 |
| B6 CD45.1 | Charles River Laboratories | Cat# CRL:564, RRID: IMSR_CRL:564 |
| B6 Thy1.1 | The Jackson Laboratory | Cat# JAX:000406, RRID: IMSR_JAX:000406 |
| B6 Cg-Thy1a/Cy Tg(TcraTcrb)8Rest/J | The Jackson Laboratory | Cat# JAX:005023, RRID: IMSR_JAX:005023 |
| Kaede B6 | Dr. M. Tomura, RIKEN & Dr. A. Luster, Harvard University | Catalog# RBRC05737, RRID: IMSR_RBRC05737 |
| B6 Ifnar1 KO | The Jackson Laboratory | Cat# 032045-JAX, RRID: MMRRC_032045-JAX |
| Mx1 GFP B6 | The Jackson Laboratory | Cat# 033219; RRID: IMSR_JAX:033219 |
| **Software and Algorithms** | | |
| FlowJo v10 | Treestar Inc | RRID: SCR_008520 |
| Prism 8 | Graphpad Inc | RRID: SCR_002798 |
| Seurat v3.0 | https://satijalab.org/seurat/get_started.html | RRID: SCR_016341 |
| Cell Ranger | 10X Genomics | RRID: SCR_017344 |
| RStudio | N/A | https://www.rstudio.com/ |
| scRepertoire | N/A | Borcherding et al. 2020, https://ncborcherding.github.io/vignettes/vignette.html#1_Introduction |

**Table S2: Reagents and Resources**

| REAGENT or RESOURCE | SOURCE | IDENTIFIER |
|---|---|---|
| **Antibodies for Flow Cytometry/Depletion and Neutralizion** | | |
| PE/Cy7 anti-mouse CD69 | Biolegend | Cat# 104512; RRID: AB_493564 |
| BV510 anti-mouse CD62L | Biolegend | Cat# 104441; RRID: AB_2561537 |
| APC anti-mouse/human CD44 | Biolegend | Cat# 103012; RRID: AB_312963 |
| BV421 anti-mouse CD103 | Biolegend | Cat# 121422; RRID: AB_2562901 |
| APC/Cy7 anti-rat CD90/mouse CD90.1 (Thy-1.1) | Biolegend | Cat# 202519; RRID: AB_2201418 |
| PE/Dazzle 594 anti-mouse CD8a | Biolegend | Cat# 100762; RRID: AB_2564027 |
| PE anti-mouse IFNAR-1 | Biolegend | Cat# 127312; RRID: AB_2248800 |
| APC anti-mouse CD45.1 | eBioscience | Cat# 17-0453-82; RRID: AB_469398 |
| PE anti-mouse/human CD44 | Biolegend | Cat# 103008; RRID: AB_312959 |
| BV711 anti-mouse CD186 (CXCR6) | Biolegend | Cat# 151111; RRID: AB_2721558 |
| FITC anti-mose CD127 (Il7ra) | eBioscience | Cat# 11-1271-82; RRID: AB_465195 |
| BV421 anti-mouse CD169 (Siglec-1) | Biolegend | Cat# 142421; RRID: AB_2734202 |
| BV421 anti-mouse/human CD45R/B220 | Biolegend | Cat# 151111; RRID: AB_2721558 |
| APC/Cyanine7 anti-mouse/human CD45R/B220 Antibody | Biolegend | Cat #103224; RRID: AB_313007 |
| AF647 anti-rat CD90/mouse CD90.1 (Thy-1.1) | Biolegend | Cat# 202508; RRID: AB_492884 |
| PE anti-mouse Ly-6D Antibody | Biolegend | Cat #138604; RRID: AB_2137349 |
| Brilliant Violet 421™ anti-mouse CD11c Antibody | Biolegend | Cat #117343; RRID: AB_2563099 |
| Brilliant Violet 510™ anti-mouse/human CD11b Antibody | Biolegend | Cat #101263; RRID: AB_2629529 |
| Depleting anti-CD4 (GK1.5) | Bioxcell | Cat# BE0003-1; RRID: AB_1107636 |
| TotalSeq™-C0301 anti-mouse Hashtag Antibody | Biolegend | Cat# 155861; RRID: AB_2800693 |

| | | |
|---|---|---|
| InVivoMAb anti-mouse CD317 (BST2, PDCA-1) | Bioxcell | Cat #BE0311; RRID: AB_2736991 |
| InVivoMAb anti-mouse Siglec-H | Bioxcell | Cat #BE0202; RRID: AB_10949014 |
| InVivoMAb anti-mouse IFNAR-1 | Bioxcell | Cat #BE0241; RRID: AB_2687723 |
| **Chemicals, Peptides, and Recombinant Proteins** | | |
| Collagenase Type IV | Worthington Biochemical | Cat# CLSS-4 |
| Dnase I | Sigma | Cat# 10104159001 |
| BSA | Sigma-Aldrich | Cat# 12659-100GM |
| Liberase | Roche | Cat# 5401020001 |
| Percoll | GE | Cat# 17089101 |
| D-Luciferin, Potassium Salt | GoldBio | Cat# LUCK-3G |
| RPMI 1640 | Corning | Cat# 10-040-CV |
| HBSS | VWR | Cat# VWRL0121-0500 |
| Live Dead Kit | ThrermoFisher | Cat# L34962 |
| 10× Single cell 3’ V2 chemistry | 10× Genomics | Cat# PN-120237 |
| Fiber Coupled Violet LED light source (Optogenetics-LED-Violet) | Prizmatix | |

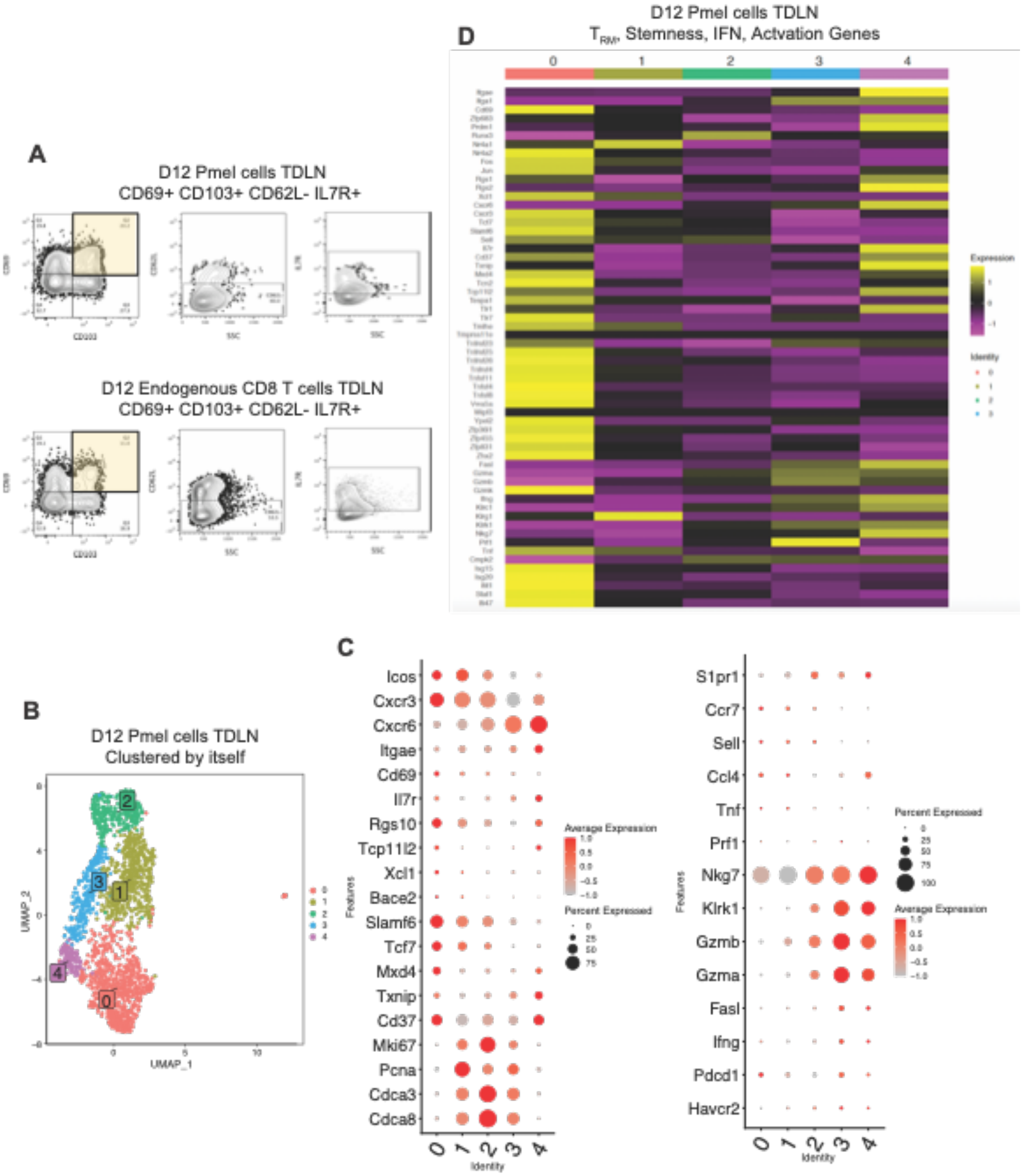

**Figure S1: scRNA sequencing analysis of pmel cells sorted from TDLNs on day 12, clustered independently.** (**A**) Flow cytometry plots showing CD69+/- CD103+/- CD62L- IL7R+ proportions on either Thy1.1+ pmel (top) or polyclonal endogenous (bottom) CD8+ T cells in d12 TDLN. (**B**) scRNA UMAP plot on d12 TDLN pmel cells. (**C**) Dotplot and (**D**) heatmap depicting gene expression on markers associated with cell cycle, stemness, tissue residency, cytotoxicity, circulation, and type 1 interferon sensing in D12 TDLN pmel cells. Flow plots are representative of 2-3 independent experiments (n=5-10; mean $\pm$ SEM).

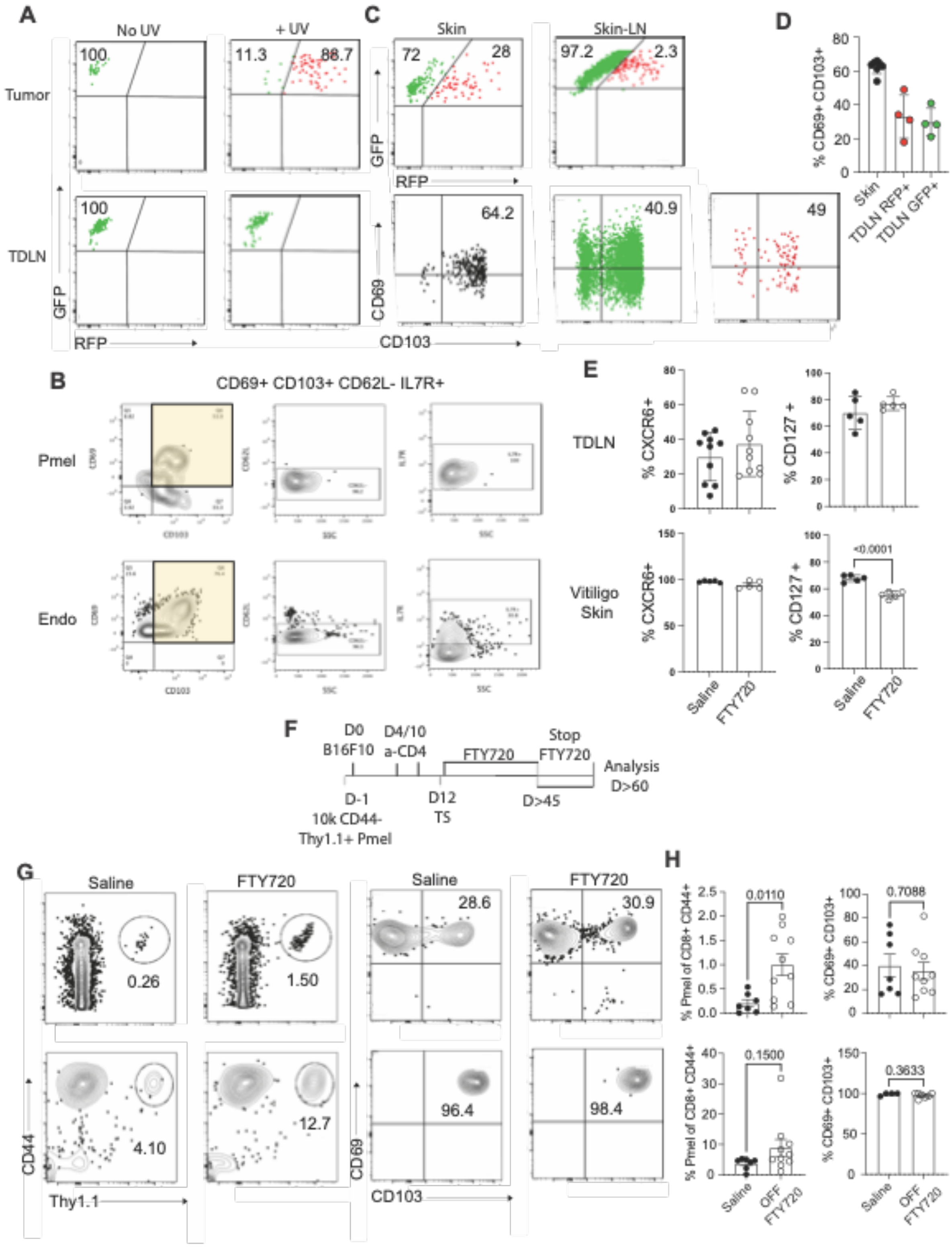
A
No UV
+ UV
Tumor
TDLN
GFP
RFP
100
11.3
88.7
100
C
Skin
Skin-LN
72
28
97.2
2.3
GFP
RFP
CD69
CD103
64.2
40.9
49
D
% CD69+ CD103+
Skin
TDLN RFP+
TDLN GFP+
B
CD69+ CD103+ CD62L- IL7R+
Pmel
Endo
E
TDLN
Vitiligo Skin
% CXCR6+
% CD127 +
<0.0001
Saline
FTY720
F
D0
B16F10
D4/10
a-CD4
FTY720
Stop FTY720
Analysis D>60
D-1
10k CD44-
Thy1.1+ Pmel
D12
TS
D>45
G
Saline
FTY720
0.26
1.50
28.6
30.9
4.10
12.7
96.4
98.4
CD44
Thy1.1
CD69
CD103
H
% Pmel of CD8+ CD44+
% CD69+ CD103+
0.0110
0.7088
0.1500
0.3633
Saline
OFF FTY720

**Figure S2: Trafficking of Trm like cells from skin to TDLN isn't required for Trm generation.** (**A**) Flow cytometry plots showing GFP/RFP proportions on Kaede Thy1.1+ pmel cells harvested 12 days post tumor inoculation from either tumor (top) or matched TDLN (bottom), without (left) or after 5 minutes (right) of UV light exposure. (**B**) Flow cytometry plots showing GFP/RFP proportions on Kaede Thy1.1+ pmel cells harvested 3 days post tumor surgery (d15) from either skin (**Top-left**) or matched TDLN (**Top-right**), after 10 minutes of UV light exposure to skin on d12-14. Bottom panel shows proportions of $CD69^{+/-}CD103^{+/-}$ gated on Thy1.1+ pmel cells. (**C**) Quantification of proportion of $CD69^{+}CD103^{+}$ out of Thy1.1+ pmel cells in skin and TDLN. (**D**) Flow cytometry plots showing CD69+/- CD103+/- CD62L- IL7R+ proportions on either Thy1.1+ pmel or endogenous cells on d12 tumor adjacent skin. (**E**) Percentage of CXCR6+ (**Left**) and CD127+ (**Right**) pmel cells from the same mice treated with FTY720 (**Fig 2J-M**). (**F**) Schematic of long term FTY720 treatment followed by a 2-week recovery period before harvest. (**G-H**) Proportions of CD44+ Thy1.1+ pmel and their phenotype (CD69/CD103) two weeks after long term FTY720 treatment was stopped. Data is representative of 2 independent experiments (**A, B, E, F, G, H**), n = 5-10; means $\pm$ SEM, or one experiment (**C, D**); n=5; mean $\pm$ S.D.

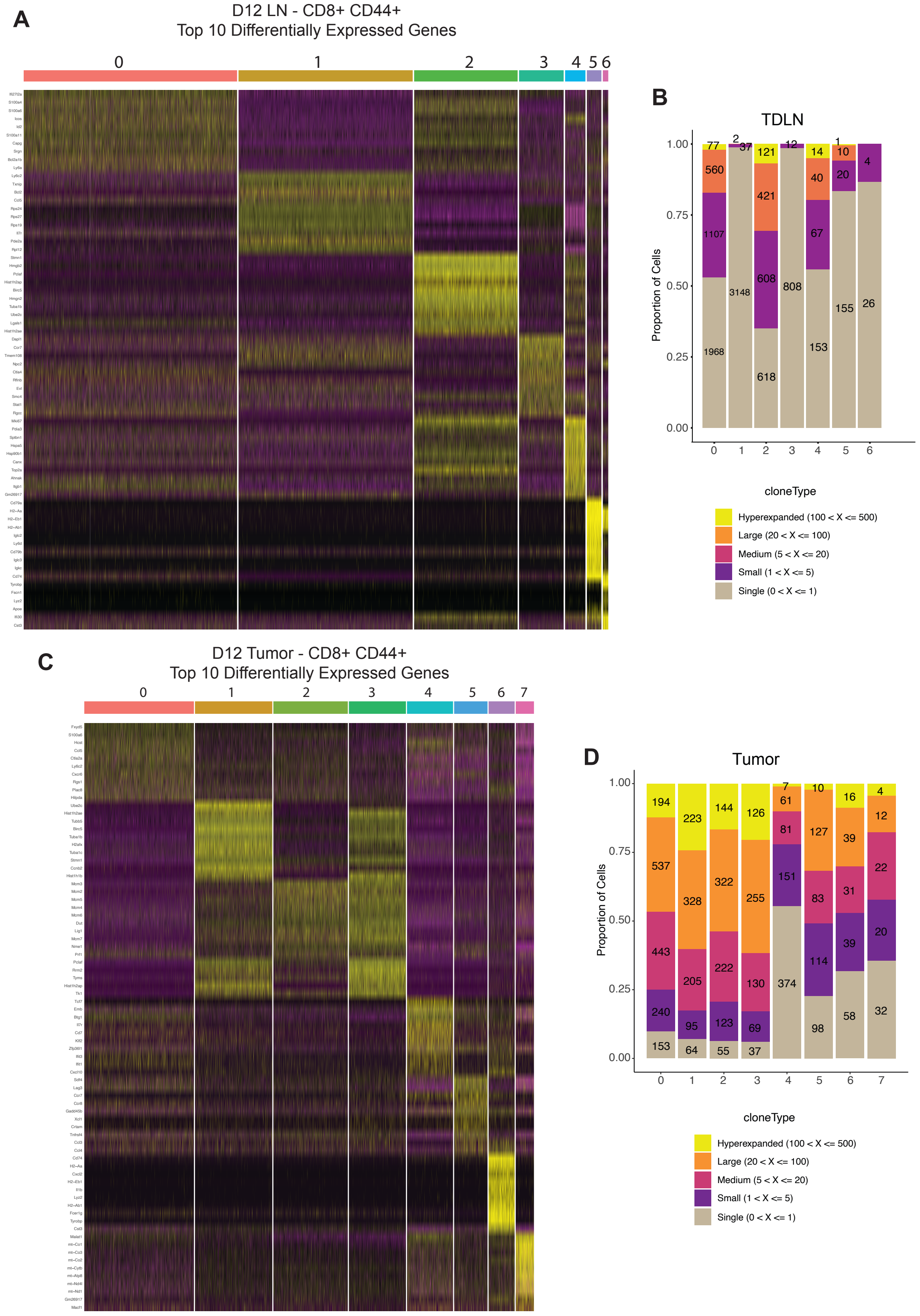
A
D12 LN - CD8+ CD44+
Top 10 Differentially Expressed Genes
0
1
2
3
4
5
6
B
TDLN
Proportion of Cells
1.00
0.75
0.50
0.25
0.00
77
560
1107
1968
2
37
3148
121
421
608
618
12
808
14
40
67
153
1
10
20
155
4
26
0
1
2
3
4
5
6
cloneType
Hyperexpanded (100 < X <= 500)
Large (20 < X <= 100)
Medium (5 < X <= 20)
Small (1 < X <= 5)
Single (0 < X <= 1)
C
D12 Tumor - CD8+ CD44+
Top 10 Differentially Expressed Genes
0
1
2
3
4
5
6
7
D
Tumor
Proportion of Cells
1.00
0.75
0.50
0.25
0.00
194
537
443
240
153
223
328
205
95
64
144
322
222
123
55
126
255
130
69
37
7
61
81
151
374
10
127
83
114
98
16
39
31
39
58
4
12
22
20
32
0
1
2
3
4
5
6
7
cloneType
Hyperexpanded (100 < X <= 500)
Large (20 < X <= 100)
Medium (5 < X <= 20)
Small (1 < X <= 5)
Single (0 < X <= 1)

**Figure S3: Trm-like cells in TDLNs acquire a distinct transcriptional signature as compared with to their clonal counterparts in tumors.** (**A-C**) Heatmap of top 10 differentially expressed genes per cluster calculated using Seurat FindMarker function from scRNA sequencing analysis on CD44+ CD8 T cells harvested from d12 TDLN and tumor. (**B-D**) scTCR sequencing analysis - Bar plot displaying clonal occupancy and expansion of CD44+ CD8 T cells harvested from d12 TDLN and d12 tumor, n=3.

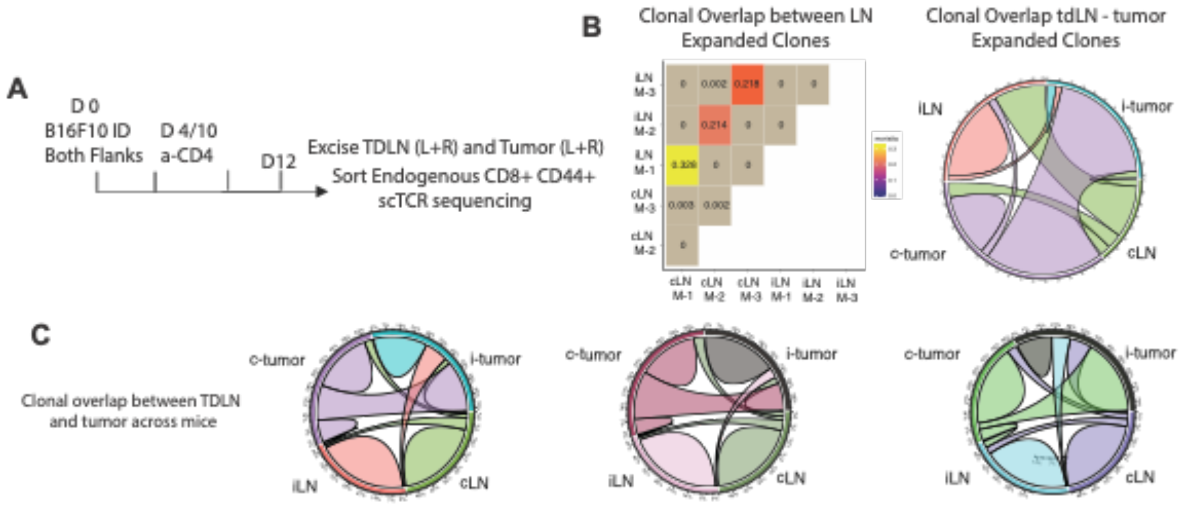

**Figure S4: TDLN excision doesn't impair clonal equilibration between tissues and Trm formation.** (**A**) Schematic showing TDLN and tumor excision (Ipsilateral and contralateral) on d12 to assess clonal overlap using scTCR sequencing on sorted CD44+ CD8 T cells. (**B-C**) Matrix and circular plot showing clonal overlap across tissues, n=3.

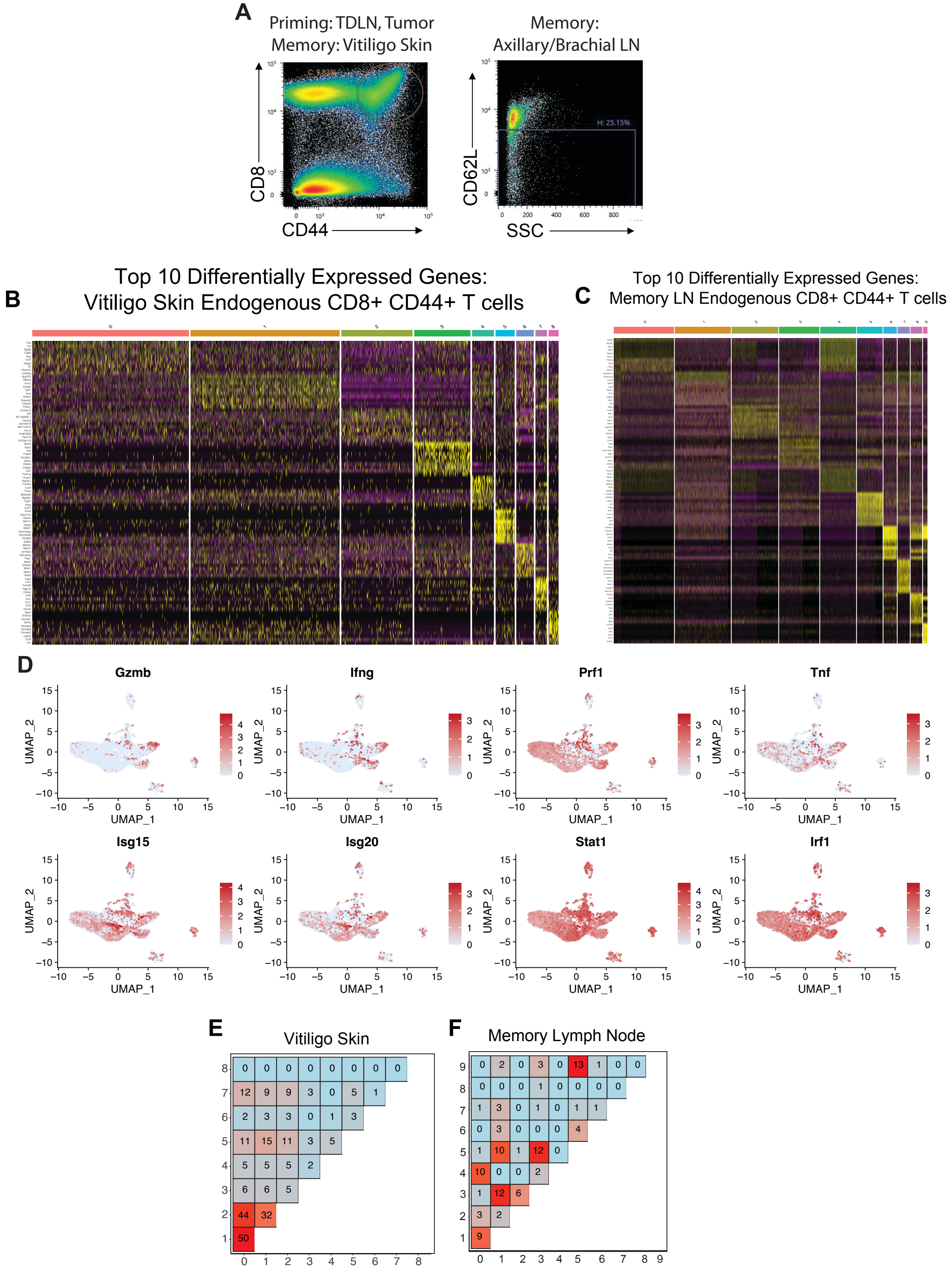
A
Priming: TDLN, Tumor
Memory: Vitiligo Skin
CD8
CD44
Memory:
Axillary/Brachial LN
CD62L
SSC
H: 25.15%
B
Top 10 Differentially Expressed Genes:
Vitiligo Skin Endogenous CD8+ CD44+ T cells
C
Top 10 Differentially Expressed Genes:
Memory LN Endogenous CD8+ CD44+ T cells
D
Gzmb
Ifng
Prf1
Tnf
Isg15
Isg20
Stat1
Irf1
UMAP_1
UMAP_2
E
Vitiligo Skin
F
Memory Lymph Node

**Figure S5: Tissue specific transcriptional signatures of CD8+ T cells in vitiligo skin and memory LN.** (**A**) Flow cytometry sorting plots of CD8 T cells sorted from d12 TDLN, tumor and d45 vitiligo skin as CD44+ and memory d45 LN as CD44+ CD62L$^{lo/int}$. (**B-C**) Top 10 differentially expressed genes per cluster calculated using Seurat FindMarker function for CD44+ CD8 T cells from vitiligo skin (**B**) or memory LN (**C**) of mice in figure 4A. (**D**) Feature plots of selected genes from scRNA sequencing on memory LN CD8 T cells from mice in figure 4A. (**E, F**) Matrix showing clonal overlap across scRNA UMAP clusters from vitiligo skin (**E**) and memory LN (**F**), n=3.

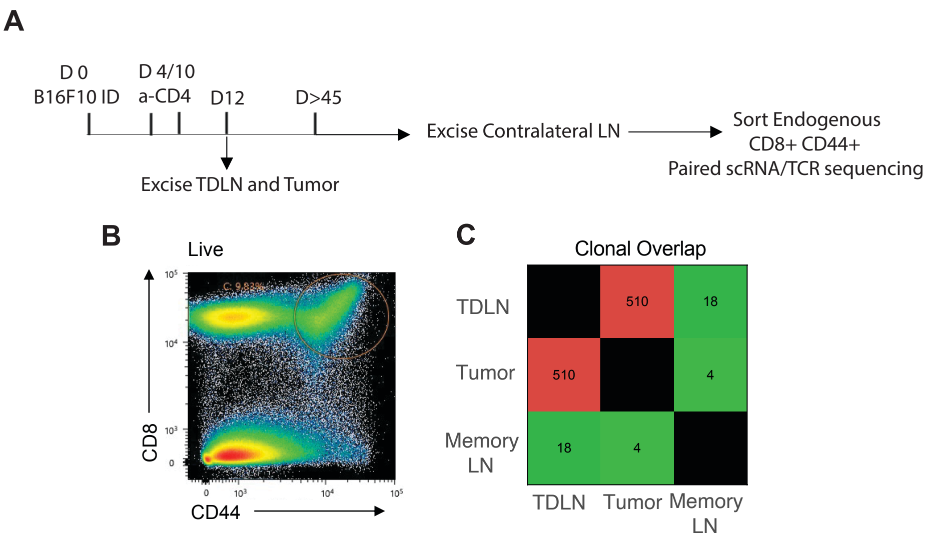

**Figure S6: Precursor T cell clonotypes from TDLNs have low clonal overlap with memory T cell populations that are not pre-enriched for Trm cells.** (**A**) Schematic showing B16 melanoma neoadjuvant treatment model. (**B**) CD44+ CD8 T cells were sorted from TDLN and tumor on d12 followed by tumor surgery and vitiligo establishment. At memory (d>45), contralateral LN was excised. Sorted cells from all indicated tissues at priming and memory time point were submitted for paired scTCR/RNA sequencing, n=3. (**C**) Matrix showing clonal overlap across tissues.

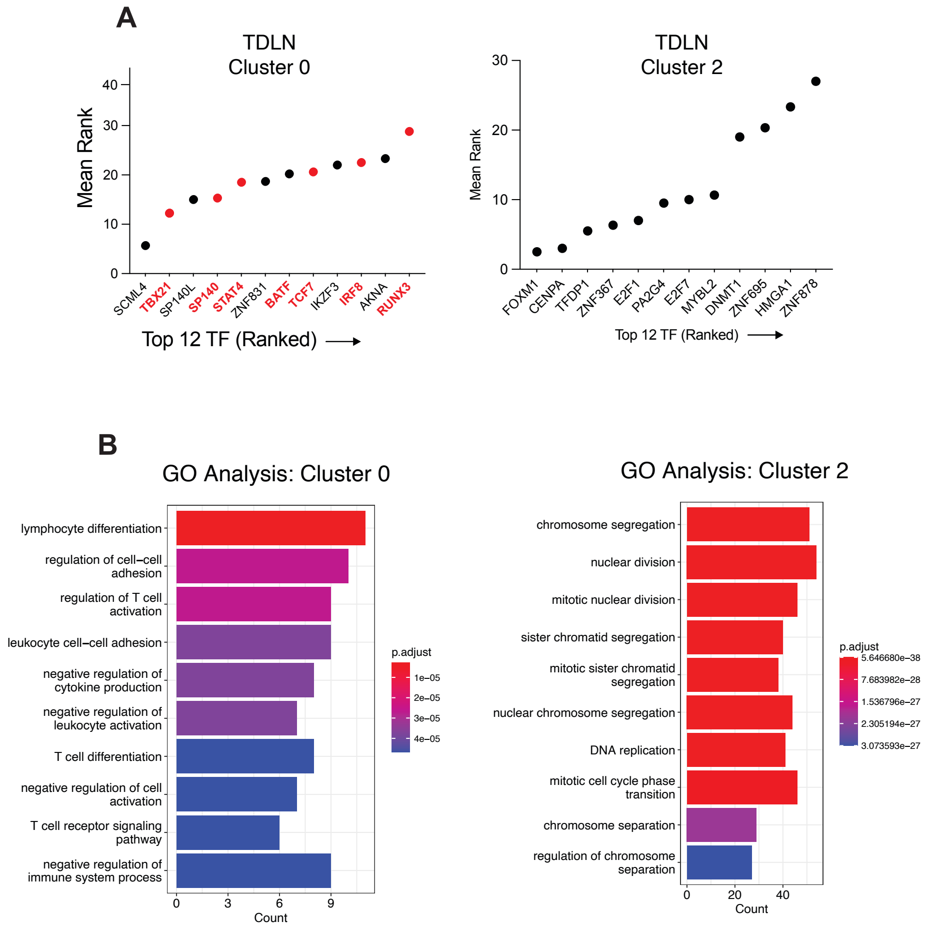

**Figure S7: Transcription factor and gene ontology analysis of TDLN_C0 (pre-Trm) and TDLN_C2.** (**A**) Transcriptional factor enrichment analysis using ChEA3 pipeline on differentially expressed genes from cluster 0 (left) and cluster 2 (right). (**B**) Gene Ontology pathway analysis using differentially expressed genes in cluster 0 (left) vs. cluster 2 (right) (**Figure 3B**), n=3.

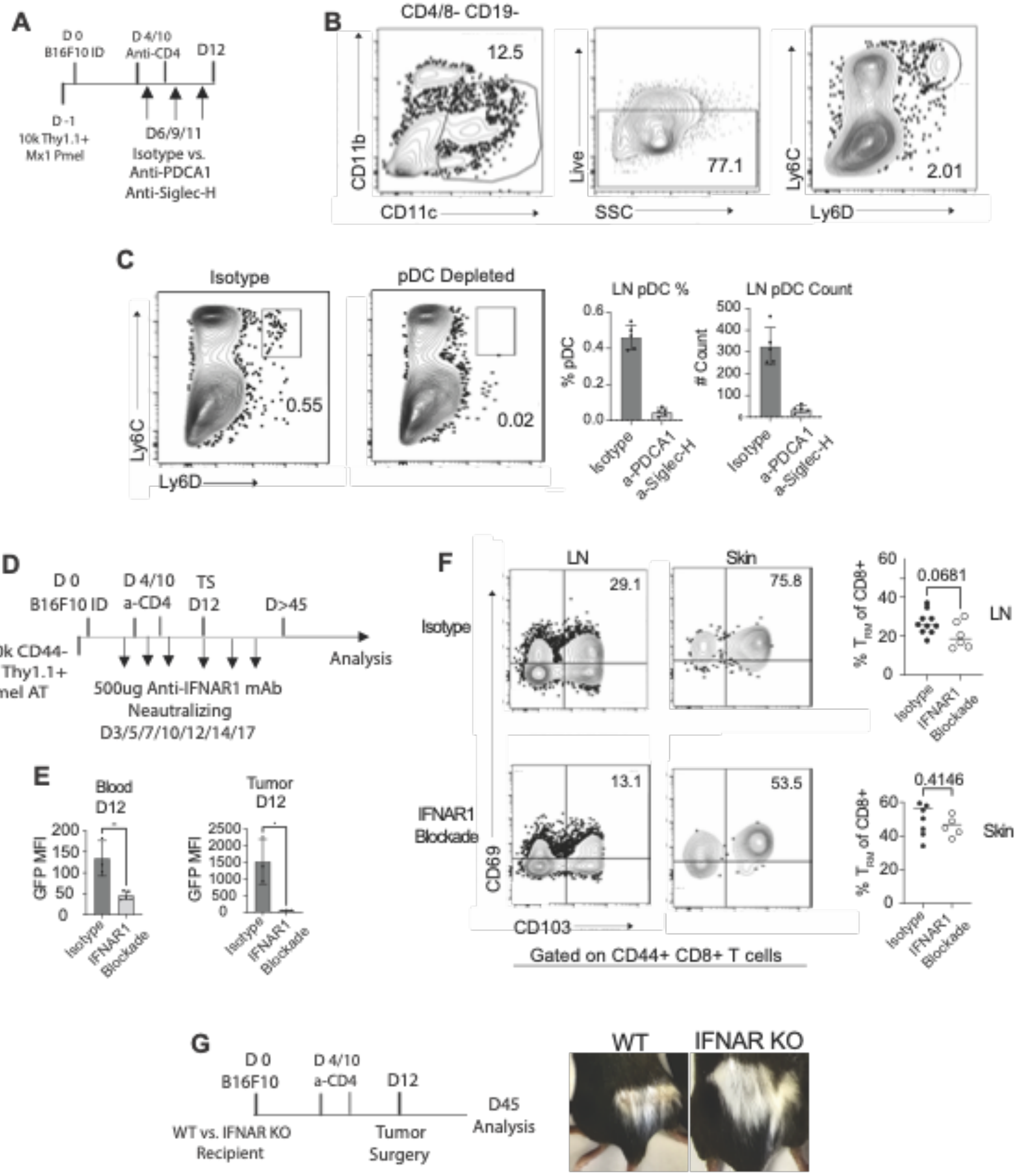

A
D 0
B16F10 ID
D 4/10
Anti-CD4
D12
D -1
10k Thy1.1+
Mx1 Pmel
D6/9/11
Isotype vs.
Anti-PDCA1
Anti-Siglec-H
B
CD4/8- CD19-
12.5
CD11b
CD11c
Live
SSC
77.1
Ly6C
Ly6D
2.01
C
Isotype
pDC Depleted
0.55
0.02
Ly6C
Ly6D
LN pDC %
% pDC
LN pDC Count
### Count
Isotype
a-PDCA1
a-Siglec-H
D
D 0
B16F10 ID
D 4/10
a-CD4
TS
D12
D>45
10k CD44-
Mx1 Thy1.1+
Pmel AT
Analysis
500ug Anti-IFNAR1 mAb
Neautralizing
D3/5/7/10/12/14/17
E
Blood
D12
Tumor
D12
GFP MFI
Isotype
IFNAR1
Blockade
F
LN
Skin
Isotype
29.1
75.8
IFNAR1
Blockade
13.1
53.5
CD69
CD103
Gated on CD44+ CD8+ T cells
0.0681
% T_RM of CD8+
LN
0.4146
Skin
Isotype
IFNAR1
Blockade
G
D 0
B16F10
D 4/10
a-CD4
D12
D45
Analysis
WT vs. IFNAR KO
Recipient
Tumor
Surgery
WT
IFNAR KO

**Figure S8: Type 1 interferon sensing imparts early Trm potential in TDLNs, and its inhibition with monoclonal antibody blockade impairs LN Trm formation.** (**A**) Schematic describing pDC depletion strategy during the priming phase using anti-PDCA1 and anti-Siglec-H depletion or isotype monoclonal antibody (mAb). (**B**) Gating strategy to identify plasmacytoid dendritic cells (pDC) (*Live* $CD4^{-}$ $CD8^{-}$ $CD19^{-}$ $CD11B^{int}$ $CD11C^{+}$ $LY6C^{+}$ LY6D. (**C**) Flow cytometry plots (left) and quantification (right) showing pDC depletion in TDLN on d12 post tumor inoculation. (**D**) Schematic describing Ifnar1 neutralization treatment using anti-Ifnar1 or isotype mAb. (**E**) GFP mean fluorescence intensity on Mx1 GFP pmel cells to confirm Ifnar1 neutralization in blood and tumor of mice that went on to develop vitiligo from panel G. (**F**) Proportion of polyclonal CD69+ CD103+ out of CD8+ CD44+ in LN and vitiligo skin of mice that received either Ifnar1 neutralizing or isotype mAb as indicated in panel D. (**G**) Vitiligo development of WT vs. IFNAR KO mice from Fig 6G. Symbols represent individual mice; horizontal lines depict means; significance was determined by paired unpaired t test, with n.s. (non-significant) denoting $p > 0.05$. Data is representative of two independent experiments, n=6-10; mean $\pm$ SEM.
